# Pseudomonads coordinate innate defense against viruses and bacteria with a single regulatory system

**DOI:** 10.1101/2025.02.26.640152

**Authors:** David M. Brinkley, Savannah K. Bertolli, Larry A. Gallagher, Yongjun Tan, Minola Motha de Silva, Ainsley Brockman, Dapeng Zhang, S. Brook Peterson, Joseph D. Mougous

## Abstract

Bacterial cells live under the constant existential threats imposed by other bacteria and viruses. Their mechanisms for contending with these threats are well documented; however, the regulation of these diverse defense elements remains poorly understood. Here we show that bacteria can mount a genome-wide, coordinated, and highly effective immune response against bacterial and viral threats using a single regulatory pathway. Bioinformatic analyses revealed that *Pseudomonas* species broadly possess a specialized form of the Gac/Rsm regulatory pathway (GRP), which our prior work in *Pseudomonas aeruginosa* implicated in activating interbacterial antagonism defense mechanisms in response to neighbor cell death. Proteomic studies comparing GRP-activated and -inactivated strains derived from diverse *Pseudomonas* species showed that the pathway regulates a large and variable suite of factors implicated in defense against both bacterial and phage threats. Focusing on *P. protegens,* we identify profound phenotypic consequences of these factors against multiple forms of bacterial antagonism and several phage. Together, our results reveal that bacteria, like more complex organisms, couple danger sensing to the activation of an immune system with antibacterial and antiviral arms.

## Introduction

For most bacteria, survival is dependent on the ability to withstand myriad biotic threats, including antagonism from phage and other bacteria. The selection imposed by these threats has led to the evolution of diverse and widespread defensive mechanisms, exemplified by the many phage defense pathways uncovered in recent years^1^. Though mechanistically distinct, these pathways employ common strategies, including blocking phage attachment, inhibiting the intracellular life cycle of the phage, or trigging programmed cell death^2^.

Interbacterial antagonism can take many forms, ranging from the production and delivery of overtly toxic molecules to more passive competition strategies that restrict nutrients^3^. Correspondingly, defense against interbacterial antagonism can present as a diffuse collection of cellular functions. However, recent work studying discrete and mechanistically defined toxin delivery systems, in particular the type VI secretion system (T6SS), has led to the identification of factors with specialized functions in defense against interbacterial antagonism. These include the production of immunity proteins that inhibit the activity of incoming toxins, extracellular barriers that block toxin delivery, pathways that repair cellular damage caused by antagonism, and counter-attack strategies^4-7^.

Defense factors are costly and the threats they protect against, though omnipresent, occur inconsistently; therefore, it stands to reason that cells might tightly regulate defense factor production^8^. The regulation of phage defense systems is perhaps most extensively documented at the posttranslational level. One common mechanism employs cyclic nucleotides, which are generated upon detection of infection or by active defense systems themselves, and then activate assorted responsive effector proteins^9^. Relatively few transcriptional and posttranscriptional mechanisms of phage defense system regulation have been identified, and it is generally thought that activation at the level of machinery biogenesis fails to keep pace with the rapid time scale of infection by many phage^10^. Indeed, quorum sensing systems are the most widely characterized class of transcriptional regulators for phage defense systems^11,12^. By virtue of extrinsic sensing of cell density, these systems permit anticipatory behavior that links high cell density to an increased likelihood of encountering a phage. A second factor driving transcriptional regulation of phage defense is the deleterious nature of many defense mechanisms for the infected cell. WYL-domain proteins are transcriptional repressors that maintain low levels of expression of several cell death-inducing phage defense pathways^13-15^. Although these proteins contain a putative ligand binding domain predicted to interact with nucleic acids to facilitate activation of the defense genes they regulate, the basal level of expression found in uninfected cells appears to be sufficient to mediate phage resistance.

Reflecting their multifaceted nature, the regulation of interbacterial antagonism defense factors is complex. Toxin delivery systems and the effectors they transport can disrupt cell wall integrity, and studies in *V. cholerae* and *E. coli* suggest that envelope stress sensing pathways could be important transcriptional activators of antagonism defense genes^16,17^. Like phage infection, certain types of interbacterial antagonism can occur on a time scale too rapid for antagonized cells to respond directly. Therefore, it is not surprising that quorum sensing is also a frequently cited regulator of interbacterial defense functions^3^. Quorum sensing can activate the production of extracellular structures that serve as barriers, and it also generally promotes the production of antimicrobials^18^. However, because quorum sensing responds to cell density rather than damage, it is an imperfect predictor of danger.

Given that common signals may predict a multitude of threats, one can envision that coordinated regulation of defense factors would be prevalent. Such a strategy is suggested by the tendency of defense genes to aggregate, such as in mobile elements containing both phage defense gene clusters and T6SS toxin genes^19^. However, outside of these situations, little is known about how genetically unlinked defense pathways are co-regulated. In *P. aeruginosa*, we previously reported the discovery of a pathway, termed the *P. aeruginosa* response to antagonism (PARA), which coordinates the induction of a broad array of interbacterial antagonism defense mechanisms in response to kin cell lysate. Together, these co-regulated factors provide multiple logs of fitness to populations confronted with highly antagonistic competitors (Figure 1A)^5,20^. PARA is mediated by the Gac/Rsm posttranscriptional regulatory program, a pathway conserved across γ-proteobacteria and reported to regulate a wide range of traits^21-23^. At the core of this pathway is the sensor kinase GacS, which phosphorylates the response regulator GacA^23^. Phosphorylated GacA activates the transcription of small RNAs, which sequester the RNA binding protein RsmA (and orthologs) from target mRNAs, thereby increasing translation. Altogether, the Gac/Rsm pathway (GRP) controls expression levels of more than 200 proteins in *P. aeruginosa*^24^. GRP targets in this species relevant to antagonism defense include the components of an antibacterial T6SS, three interbacterial antagonism defense gene clusters (antagonism resistance clusters 1-3; Arc1-3), and the extracellular polysaccharide Psl^5,24,25^.

**Figure 1.**
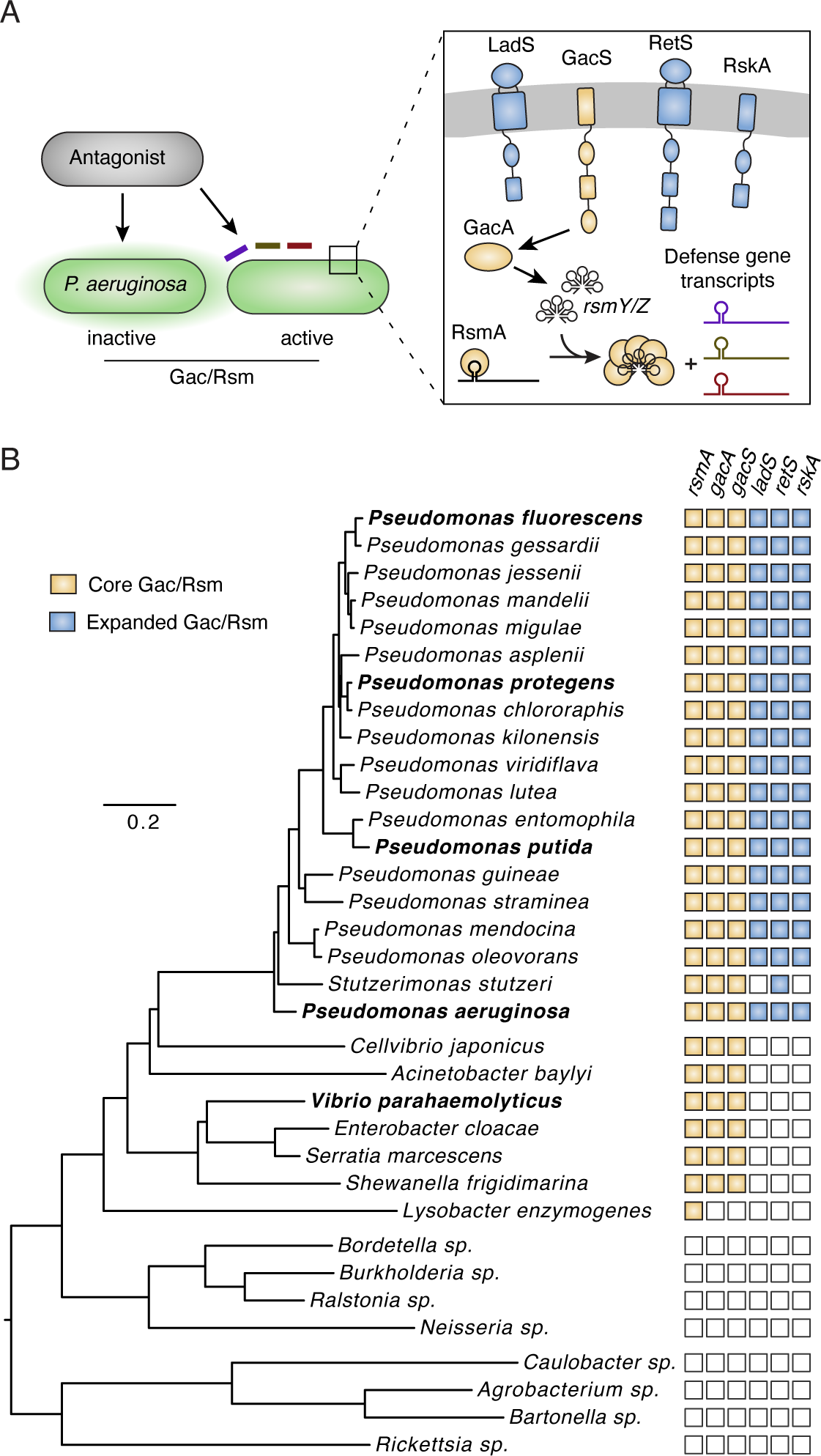
*Pseudomonas* species encode an expanded Gac/Rsm pathway. A) Diagram depicting PARA (left) and the different configurations of the Gac/Rsm pathway described in the literature (right). Note that only one RsmA homolog and two small RNA molecules (RsmY and RsmZ) are depicted for simplicity; the number of homologs of these components varies across genomes. B) Concatenated marker gene-based phylogeny of *Pseudomonas* species and representatives from other γ-proteobacterial lineages indicating GRP components identified for each in our analyses.

In this study, we show that the GRP of diverse *Pseudomonas* species regulates a variable suite of defenses against both interbacterial antagonism and phage infection. We experimentally demonstrate the central importance of the GRP in mediating defense against multiple forms of interbacterial antagonism and show that activation of the pathway provides potent defense against infection by phage. Our results support a model in which the GRP serves as a highly effective immune response for countering both bacterial and viral threats across *Pseudomonas* species.

## Results

### Pseudomonas species possess a unique form of the Gac/Rsm pathway

The Gac/Rsm signaling pathway (GRP) is widely distributed amongst γ-proteobacteria^23^. Despite some level of characterization in many of these organisms, only in *P. aeruginosa* has the pathway been implicated in defense against interbacterial antagonism^5,20,26^. Notably, the GRP of *P. aeruginosa* is augmented by the sensor kinases RetS, LadS, and RskA. We previously showed that RetS repression of GacS modulates the activity of the pathway during antagonism, and both LadS and RskA counter the activity of RetS (Figure 1A)^20,24,27,28^. Given these observations, we sought to define the distribution of both the core components of the Gac/Rsm signaling system and the modulatory sensor kinases of *P. aeruginosa*.

We identified homologs of GacS, GacA, RsmA, RetS, LadS, and RskA in representative genomes across the major proteobacteria lineages. We then subjected homologs to phylogenetic analysis, allowing us to define candidate orthologous sequences across genomes (Figure S1). Finally, we compared the domain architecture of candidates to the corresponding *P. aeruginosa* proteins, permitting a confident determination of orthologs. As previously reported, we found the core Gac/Rsm proteins GacS, GacA, and RsmA are widely distributed across γ-proteobacteria and absent outside of this class (Figure 1B)^29^. By contrast, our analysis showed that the accessory kinases RetS, LadS, and RskA are strictly confined to organisms belonging to the genus *Pseudomonas*. Although *retS*, *ladS* and *rskA* are not genetically linked, with only one exception (*P. stutzeri*), the genes universally co-occurred within the *Pseudomonas* spp we queried. Together, these data show that *Pseudomonas* spp broadly possess a unique version of the Gac/Rsm pathway that is potentially capable of mediating sensing of and response to antagonism.

### Mass spectrometry of regulatory mutants sensitively defines the GRP regulon

We previously demonstrated that the critical role of the *P. aeruginosa* GRP in defense against interbacterial antagonism derives from the cumulative effect of multiple individual factors under its control^5^. Here, we sought to determine how the GRP regulon varies across a phylogenetically diverse cross-section of Pseudomonas species. We hypothesized that the identity of factors under GRP control in these species would provide general insights into its function beyond *P. aeruginosa*.

To our knowledge, there has not been an effort to apply consistent methodology to experimentally define the GRP regulon across multiple species of *Pseudomonas*. However, groups have undertaken various approaches to define the regulon in individual species. These include bioinformatic analyses to predict transcripts bound by RsmA, RsmA ChIP-seq and related methods, and transcriptome measurements ^24,30-35^. As a post-transcriptional regulatory pathway, the GRP regulon is arguably most appropriately characterized at the protein level. Only a small number of studies have reported such data, and these compared protein expression between GRP-inactive (e.g. Δ*gacS*) and wild-type strains^26,33^. However, our prior data suggest that activation of the pathway under *in vitro* growth conditions requires an antagonizing organism or deletion of a negative regulator (e.g. Δ*retS*)^20,36^. We therefore reasoned that in the absence of antagonism, GRP-regulated factors might be most sensitively identified in a given *Pseudomonas* species by comparing the proteome of a GRP-inactivated strain to that of one in which a negative regulator of the pathway is removed. To test this, we compared whole cell proteomes of *P. aeruginosa* wild-type, Δ*gacS*, and Δ*retS* strains using principal component analysis. Along PC1 (42% of variance), Δ*retS* proteomes diverged substantially from wild-type and displayed a greater distance from Δ*gacS* than did the wild-type (Figure 2A). We conclude that GRP regulons of *Pseudomonas* spp can be most comprehensively defined by comparing protein abundances between Δ*retS* and Δ*gacS* genetic backgrounds.

**Figure 2.**
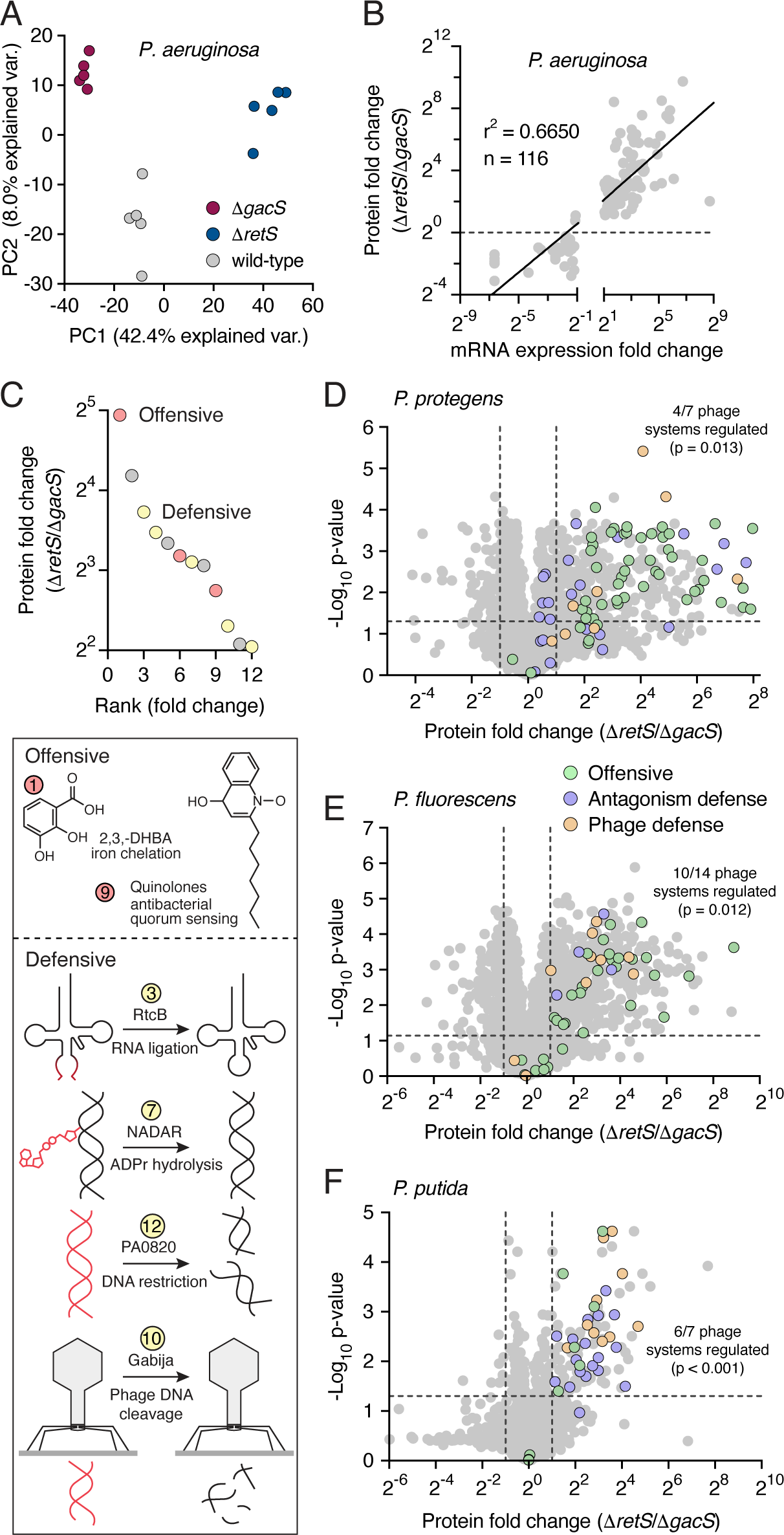
Proteomic analysis of the GRP regulon of diverse *Pseudomonas* species. A) Principal component analysis of MS-derived whole-cell proteomes (n=5) from the indicated strains of *P. aeruginosa.* B) Comparison of the fold change in abundance between *P. aeruginosa* Δ*retS* and Δ*gacS* strains for GRP targets as measured by transcript (X-axis, derived from Goodman *et al.*^24^) or protein abundance (this study). C) Plot showing most differential previously undetected GRP targets in *P. aeruginosa* by rank fold change (left) and schematic illustrating antimicrobial and defensive functions associated with a subset of these targets. D-F) Volcano plots comparing differences in abundance of proteins detected by MS for Δ*retS* and Δ*gacS* strains of *P. protegens* (C), *P. fluorescens* (D) and *P. protegens.* Average fold-change and p-values were calculated from 5 biological replicates.

We began our investigation of the GRP regulon in *Pseudomonas* spp. by benchmarking our *P. aeruginosa* dataset against two prior transcriptomic studies^24,32^. A strong correlation in abundance between significant targets was observed with both datasets, and consistent with expectations, a transcriptomic study comparing GRP-activated and -inactivated backgrounds more closely resembled our findings than did the study employing a wild-type reference (Figure 2B, Figure S2A). In total, our determination of the GRP regulon of *P. aeruginosa* supports prior data implicating the pathway in antimicrobial production and antagonism defense. Notable antimicrobial GRP targets confirmed by our data include the antibacterial H1-T6SS and 1- undecene biosynthesis proteins, whereas antagonism defense targets include Arc pathway components and Psl exopolysaccharide biosynthesis proteins^24^.

In addition to the previously identified targets in our dataset, we found 72 GRP-regulated proteins not identified in prior genome-wide studies (Supplementary Table 1). Lending credence to the validity of these proteins as bona fide components of the GRP regulon, the group includes multiple proteins defined as GRP targets in focused investigations. Among these are the stationary phase sigma factor RpoS, the quorum sensing signal synthase LasI, and genes under the control of the transcription activator BexR^37-39^. Ranking the previously undescribed GRP targets within our data by induction level revealed that a preponderance of those most strongly induced by the pathway represent additional antimicrobial and antagonism defense functions (Figure 2C). For instance, KynA, B and U generate an essential precursor of quinolone biosynthesis^40^. Several quinolones of *P. aeruginosa* are antimicrobial, and one of these, PQS (*Pseudomonas* quinolone signal), further stimulates production of the antimicrobial pyocyanin via quorum sensing^41,42,43^. We also identified the enzyme responsible for biosynthesis of the siderophore 2,3-dihydroxybenzoic acid^44^. Siderophores are well known to function in microbial competition by depleting accessible iron from neighboring microbes^45^. Previously undescribed GRP targets we identified with defensive functions include two that counteract the effects of antibacterial toxins: a homolog of RtcB that repairs RNA molecules cleaved by RNase toxins and a NADAR (NAD- and ADP-ribose-associated) family protein, members of which remove ADP ribose moieties installed by ADP-ribosylating toxins on nucleic acids^46-49^. Other defensive factors include a predicted restriction endonuclease (PA0820) and a homolog of the DNA replication inhibitor CspD, which promotes persister cell formation^50^. Notably, we identified the phage defense system Gabija among the most strongly induced, previously undescribed GRP targets of *P. aeruginosa*. This raised the intriguing possibility that the defensive function the GRP could extend to viruses. Together, these results validate our proteomics-based approach and significantly expand our understanding of the GRP regulon in *P. aeruginosa*. Furthermore, they strongly support the contention that the GRP of this bacterium controls factors that constitute a multifaceted response to a range of antagonistic threats.

### Interbacterial antagonism and phage defense mechanisms dominate the GRP regulon in diverse *Pseudomonas* species

Given the insights garnered by our MS-based interrogation of the GRP regulon of *P. aeruginosa*, we applied the strategy to a diverse cross-section of other Pseudomonads. We selected *P. protegens*, *P. fluorescens*, and *P. putida* for these studies, primarily owing to their phylogenetic divergence from each other and from *P. aeruginosa*; however, these species also inhabit a range of distinct ecological niches, suggesting exposure to unique sets of biotic threats^51,52^. Application of our proteomics-based approach to these species revealed extensive proteomic remodeling by the GRP pathway, similar in magnitude to that observed in *P. aeruginosa,* as well as substantial differences between the proteomes of each species (Figure 2D- F and Supplementary Figure 2B-D). Relatively few proteins are under GRP control in all four species. Notably, all but one of these shared GRP targets represent functions related to interbacterial antagonism or defense. These include components of the H1-T6SS and the outer-membrane stress resistance protein LptF ^53^.

Beyond conserved GRP targets, we found extensive variability among regulated targets across the four species investigated. However, as in *P. aeruginosa,* the GRP regulon of each species encompassed a preponderance of functions related to antimicrobial production or defense against antagonism. For instance, the GRP regulons of both *P. protegens* and *P. fluorescens* include the biosynthetic machinery for production of numerous secondary metabolites, many of which have antimicrobial activity. Consistent with prior studies, we found the proteins responsible for production of 2,4-DAPG, rhizoxin, protegenin, pyoluteorin and orfamide in *P. protegens* are positively regulated by the GRP (Supplementary Table 1)^54^. Expression of a candidate orfamide biosynthetic pathway is also activated by the GRP in *P. fluorescens,* as well as proteins belonging to an uncharacterized non-ribosomal peptide synthesis (NRPS) pathway (PFLU3_17930-PFLU3_17980) and the biosynthetic pathways for phenazine and azetidomonamide. Defensive functions under GRP control in these species include production of the extracellular polysaccharides Psl and Peb in *P. protegens* and *P. putida*, respectively, multiple universal stress protein homologs in *P. putida*, and periplasmic proteins in *P. putida* and *P. protegens* with structural similarity to BepA. This protein promotes maturation of the BAM complex and degrades misfolded outer-membrane proteins during stress, promoting cell envelope integrity^55^.

Interestingly, we identified phage defense systems within the GRP regulons of all four *Pseudomonas* spp analyzed. These achieved statistically significant association with the GRP regulons of *P. protegens* (p = 0.013), *P. fluorescens* (p = 0.012), and *P. putida* (p <0.001), where 4 of 7, 10 of 14, and 6 of 7 detected phage defense systems, respectively, are GRP regulated (Figure 2D-F and Supplementary Table 3). Components of several phage defense systems, including PARIS and BstA in *P. protegens* and Shango and Kiwi in *P. fluorescens*, are above the 85^th^ percentile of GRP induction level in these bacteria. This finding, taken together with the multitude of GRP-regulated antibacterial and antifungal pathways established by our MS-based approach, strongly suggests that the GRP regulon constitutes a central immune response with arms targeting each major category of biotic threat.

### Pseudomonas sp. broadly employ Gac/Rsm in antagonism defense

The GRP regulons we defined in *P. putida*, *P. protegens*, and *P. fluorescens* are consistent with the pathway broadly mediating defense against microbial antagonism beyond *P. aeruginosa*. As a first experimental test of this, we measured the benefit of GRP activity in our panel of Pseudomonads under conditions of interbacterial antagonism by the T6SSs of *Burkholderia thailandensis* (*B. thai*) and *Enterobacter cloacae*. In competition with *B. thai*, Δ*gacS* strains of each species tested exhibited severe antagonism-dependent fitness defects (3-6 log, Figure 3A). With the exception of *P. fluorescens*, the Δ*gacS* strains of each species also displayed pronounced antagonism defense defects against *E. cloacae*. Importantly, the GRP- inactivated strains did not display a growth defect in isolation or in co-culture with antagonism- deficient competitors (Supplementary Figure 3A,B). Prior work suggests that outside of *Pseudomonas*, strains bearing GRP-inactivating mutations can exhibit marked *in vitro* growth defects^56-59^. We found an exception to this pattern is *Vibrio parahaemolyticus*, which allowed us to evaluate the contribution of the pathway to antagonism defense in a bacterium lacking the accessory kinases without this confounding variable (Supplementary Fig. 2C). Although antagonized effectively by the T6SSs of both *B. thai* and *E. cloacae*, we did not observe an antagonism-dependent fitness defect of *V. parahaemolyticus* Δ*gacA* against these competing organisms (Figure 3A, Supplementary Fig. 3D). Our experiments reveal that the specialized GRP of *Pseudomonas* is uniquely adapted for defense.

**Figure 3.**
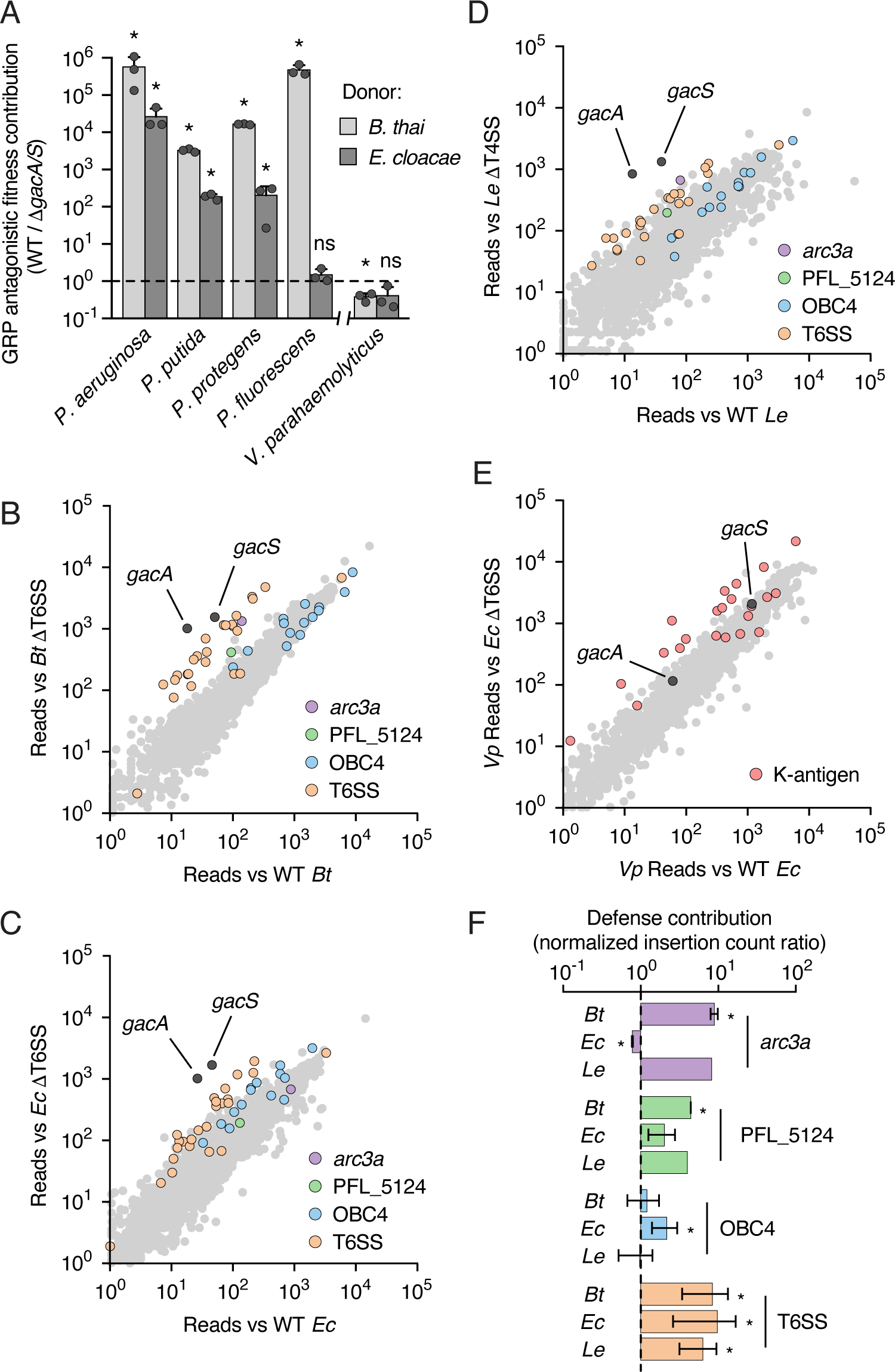
The GRP plays a general and central role in defense against interbacterial antagonism. A) Mean relative fitness (±SD) of the wild-type strain for each species during T6SS-mediated antagonism by the indicated competitors, relative to the fitness of the corresponding GRP-inactivated mutants. Asterisks indicate strain pairings in which we observe a significant antagonism-dependent fitness difference between the wild-type and Δ*gacS* strains. (Welch’s T-test, p<0.05 with BH correction for multiple comparisons, n=3). B-E) Transposon library sequencing-based comparison of the fitness contribution of individual *P. protegens* (B-D) or *V. parahemolyticus* (E) genes during growth competition with wild-type versus antagonism deficient *B. thai* (B), *E. cloacae* (C, E) or *L. enzymogenes* (D). GRP signaling components (GacA, GacS) indicated in B-E, and GRP-targets that contribute to defense against one or more antagonist are highlighted in B-D. F) Contribution of the indicated pathways to defense against different competitors. Average normalized transposon insertion count ratios (antagonism deficient competitor/WT competitor, ±SD) were obtained using all genes within a given pathway, and replicate Tn-seq datasets when available. For Arc3 and PFL_5124, a single ratio measurement is provided for defense against *L. enzymogenes*, as a single Tn-seq screen was performed with that antagonist. Asterisks indicate ratios significantly different from 1 (1-sample t-test, p<0.05)

### The GRP is the major defense determinant for *P. protegens* against multiple mechanisms of interbacterial antagonism

We next sought to contextualize the impact of GRP inactivation on defense against antagonism by comparing its contribution relative to other, potentially GRP-independent, defense mechanisms. To accomplish this in a comprehensive and unbiased manner, we conducted genome-wide transposon insertion sequencing (Tn-seq) screens wherein a *P. protegens* transposon insertion library was subjected to co-cultivation with wild-type and antagonism-deficient competitor bacteria. Given our results above, selected competitor bacteria included *B. thai*, *E. cloacae*, and their corresponding T6S-inactivated derivatives. To capture a broader range of antagonistic mechanisms, we also included *Lysobacter enzymogenes* and a derivative of this strain bearing an inactivated Xanthomonadales-like type IV secretion system (X-T4SS). Unlike conjugative T4SSs, this apparatus is specialized for the delivery of large cocktails of antibacterial proteins^60^. Notably, these toxins are distinct from those delivered by T6SSs^61^. Strikingly, *gacA* and *gacS*, showed the strongest antagonism-dependent fitness contribution across all three competitors, with average selection ratios of 178 and 188, respectively (Figure 3B-D). This finding is consistent with prior results showing that the *P. aeruginosa* GRP plays a similarly critical role in defense against T6S-based antagonism^5,20^. To compare the relative contribution of GRP and non-GRP defense mechanisms between Pseudomonads and a bacterium possessing only the basic GRP, we performed a similar selection using a transposon library of *V. parahaemolyticus* co-cultured with *E. cloacae*. Consistent with our findings using individual mutations, *V. parahaemolyticus gacA* and *gacS* mutations were not under selection during antagonism (Figure 3E). Rather, genes belonging to the K-antigen biosynthesis pathway constituted the majority of those whose function was most highly selected by antagonism. Selection of these genes suggests a defensive strategy solely reliant on blocking the T6SS via modification of the cell surface rather than a coordinated regulatory response^7,62^. Together, these genome-wide screens highlight the specialized and essential role that the extended GRP of Pseudomonads plays in antagonism defense.

In-line with the strong selection for the core GRP signaling elements in *P. protegens,* we found its GRP regulatory targets overrepresented among genes contributing to fitness during antagonism. Against all three antagonists, genes encoding essential components of the T6SS system constituted many of the most critical survival determinants (Figure 3F). Other GRP targets we identified as important for fitness during antagonism varied depending on the antagonizing organism, suggesting their functions may be specialized to provide protection from specific toxins or delivery mechanisms. For instance, we found that Arc3 provides *P. protegens* defense against *B. thai* and *L. enzymongenes,* whereas the pathway is dispensable during antagonism by *E. cloacae* (Figure 3F). Arc3 specifically grants defense against Tle3 family phospholipases, which *B. thai,* but not *E. cloacae*, possesses^5,63,64^. Based on its genome sequence, *L. enzymogenes* encodes multiple antibacterial T4SS effectors, at least one of which we found is a phospholipase that could be a Tle3 family member^65^.

Additional GRP targets we found to have antagonist-specific contributions to fitness include PFL_5124 and genes within the O-polysaccharide (O-PS) biosynthesis cluster 4 (OBC4), responsible for production of a long version of the polymer^66^. The closest structural homolog of PFL_5124 is BepA (E-value, 1.8 x 10^-7^), a periplasmic protease that, as noted above, acts to promote membrane integrity by degrading misfolded proteins that accumulate in the periplasm during stress^55^. Our Tn-seq screens indicated this protein specifically contributes to defense against *B. thai,* which encodes two T6SS effectors predicted to act in the periplasm^67^. Pairwise competition assays employing an in-frame, unmarked deletion of PFL_5124 confirmed its contribution to defense against antagonism by *B. thai* and not *E. cloacae* (Supplementary Figure 4A). Given that LPS production was not linked to the GRP prior to our proteomics analysis, we investigated the impact of pathway activation on long O-PS production. Comparison of LPS profiles of *P. protegens* Δ*gacS* and Δ*retS* strains revealed substantially less of the OBC4 polysaccharide in the GRP-inactive background (Supplementary Figure 4B,C). To further evaluate the role of the OBC4 pathway in defense, we performed growth competition assays between wild-type and ΔOBC4 strains against *E. cloacae*, *B. thai* and *L. enzymogenes*. Consistent with our Tn-seq data, OBC4 contributed to *P. protegens* fitness specifically during antagonism by *E. cloacae* (Supplementary Figure 4D).

We noted that the magnitude of the competitive defect derived from GRP-inactivating insertions is substantially greater than those that inactivate any single pathway in its regulon, and indeed greater than the predicted cumulative deficit of all the GRP-regulated pathways we hit (Figure 3F). One likely explanation for this finding is that, with many different GRP-targets acting together to promote fitness during antagonism, few individual factors make a substantial enough impact to be detected in our screens. However, we considered two additional factors that could be contributing. First, under the contact-promoting conditions of our screen, the strong fitness contribution of the T6SS could be masking other GRP-regulated defense functions. Additionally, a substantial component of the characterized GRP regulon across *Pseudomonas* species consists of antimicrobial secondary metabolite biosynthesis machineries. These pathways may well contribute to defense, but under the conditions of our screens, mutations within them would be complemented in trans by other clones in the population.

To evaluate whether the activity of the T6SS could be masking the importance of other defense pathways, we repeated our Tn-seq screen using a *P. protegens* strain lacking T6SS activity. For this experiment, we generated a transposon mutant library in *P. protegens* Δ*tssM* and selected this library, as previously, in the presence and absence of T6SS-based antagonism by *E. cloacae.* As expected for this genetic background, T6SS genes did not show selection during antagonism (Supplementary Figure 4E). On the contrary, *gacA* and *gacS* genes remained the most critical defense determinants, underscoring the importance of non-T6SS-regulated functions in defense. Further analysis of these data revealed several GRP regulon defense factors not identified in our prior screens employing the wild-type *P. protegens* background (Supplementary Figure 4F). These include *maf_1* and an uncharacterized toxin-antitoxin module (PFL_0652-0653). Maf proteins exhibit diphosphatase activity against nucleotide triphosphate and are implicated in maintaining genome integrity through degradation of non-canonical, potentially mutagenic bases^68^. Toxin-antitoxin systems are widely implicated in resistance to phage, yet to our knowledge these modules are not known to contribute to defense against interbacterial antagonism.

We next assessed the possibility that secondary metabolite production represents a second class of GRP targets important for defense that were not detected in our Tn-seq screen. Specifically, we asked whether production of the antibacterial and antifungal molecule 2,4- DAPG benefits *P. protegens* during antagonism. We inactivated the 2,4-DAPG biosynthetic pathway via in frame deletion of *phlD*, and subjected this strain to growth competition assays*. P. protegens* Δ*phlD* exhibited a significant fitness defect during antagonism with *B. thai,* reflecting the susceptibility of this species to synthetic 2,4-DAPG (Supplementary Figure 4G,H). These results support our hypothesis that antimicrobial production constitutes an important arm of the GRP-regulated immune program. Together, our Tn-seq data underscore the central importance of the GRP in coordinating a multifaceted defense against interbacterial antagonism.

### The GRP of *P. protegens* provides defense against phage

The presence of multiple phage defense pathways within the GRP regulons of diverse *Pseudomonas* species led us to hypothesize that GRP innate immune capacity extends beyond interbacterial antagonism to include a second arm dedicated to countering viral threats. To test this, we isolated lytic phages from a variety of rhizosphere soil samples using the predicted phage-sensitive strain *P. protegens* Δ*gacS* as bait. Whole genome sequencing and phylogenetic analysis revealed that our collection included six distinct, tailed double-stranded DNA phages from the class Caudoviricetes, most closely related to the *Pseudomonas* phages vB_PpuP-Luke-3 and PollyC. We then assessed plaquing efficiency of these phages on *P. protegens* GRP activated versus inactivated strains. Strikingly, we found that activation of the GRP significantly restricted the growth of each phage (Figure 4A,B).

**Figure 4.**
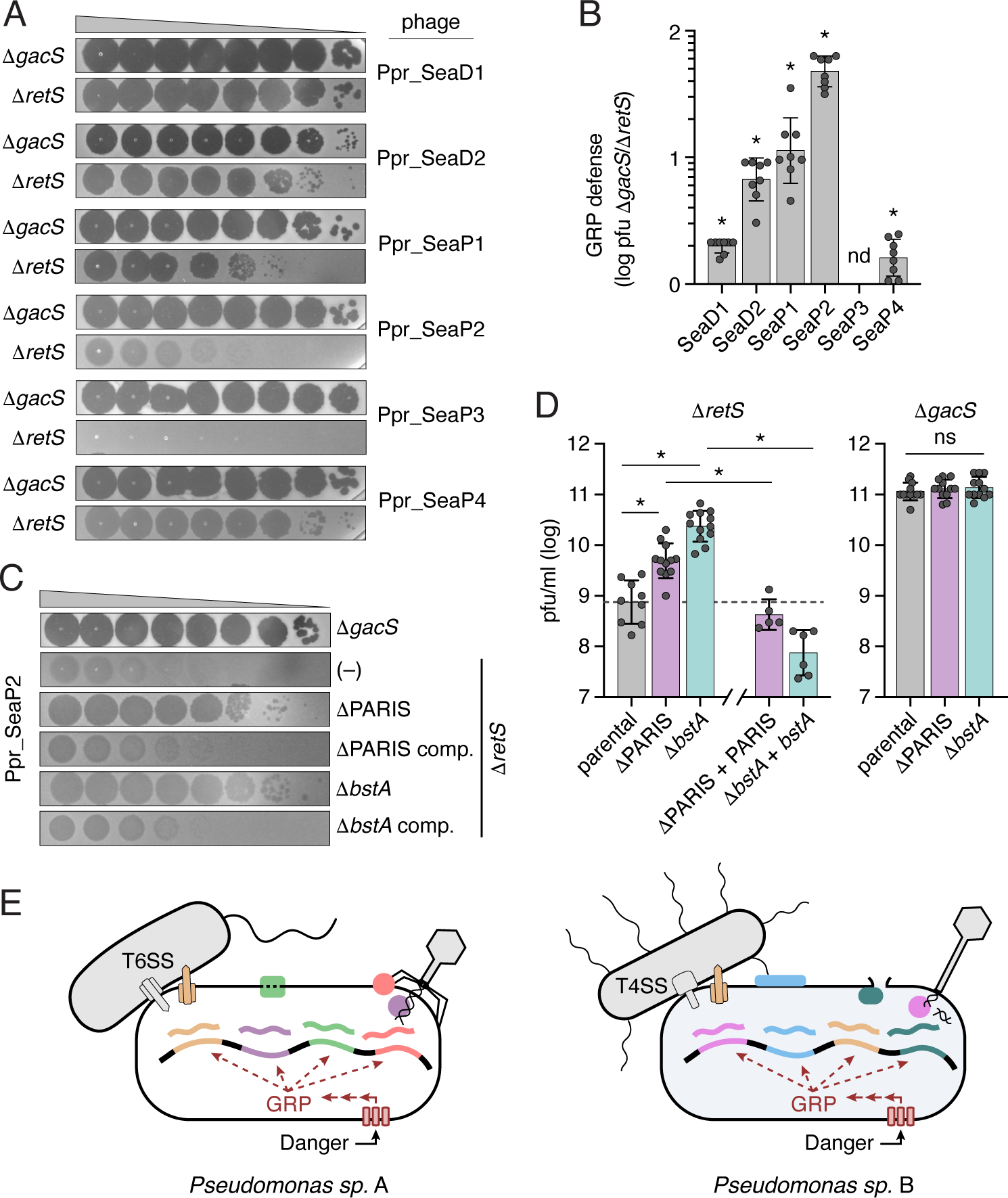
The GRP and defense factors under its control contribute to phage resistance. A) Plaquing efficiency of the indicated phages on GRP inactive (Δ*gacS*) or activated (Δ*retS*) *P. protegens.* B) Ratio of pfu obtained from the indicated phages on *P. protegens* Δ*gacS* vs Δ*retS*. Asterisks indicate ratios significantly different from one (1-sample t-test, n=8, P<0.05). ND, not determined. C) Plaquing efficiency of Ppr_SeaP2 on the indicated strains of *P. protegens.* D) Quantification of the plaquing efficiency of Ppr_SeaP2 on the indicated mutants of *P. protegens* in a GRP activated (Δ*retS*) or inactivated (Δ*gacS*) background. Asterisks indicate significant differences in PFU yield between strains (one-way ANOVA followed by Holm-Sidak post hoc test, n=5-12, p<0.05). E) Model for the role of GRP in a general coordinating immune response in *Pseudomonas* species. A common danger signal (lysate from neighboring kin cells) activates a variable suite of defenses depending on the species (indicated by different colors). These defenses protect against multiple forms of interbacterial antagonism (e.g. T4SSs and T6SSs) and infection by different phages.

The phage defense pathways we identified as most strongly induced upon GRP activation in *P. protegens* are BstA and PARIS. To determine whether the restriction of phage replication we observed upon GRP activation is attributable to these pathways, we generated in-frame deletions of each pathway in both GRP-activated and inactivated backgrounds of *P. protegens*. We found that in *P. protegens* Δ*retS*, inactivation of BstA and PARIS significantly increased plaquing efficiency by phage Ppr_SeaP2 (Figure 4C, D). Notably, the contribution of BstA and PARIS to phage resistance requires GRP activation, as inactivation of these pathways did not affect sensitivity of *P. protegens* Δ*gacS* to Ppr_SeaP2 infection (Figure 4D). Additionally, resistance to Ppr_SeaP2 is restored to the *P. protegens* Δ*retS* defense pathway mutants when complemented by the relevant pathway under the pBAD promoter at an ectopic location on the chromosome (Figure 4C, D). For the other phages tested, inactivation of these pathways did not affect the resistance conferred by RetS inactivation (Supplementary Figure 5). We speculate that for these phages, the GRP could negatively regulate a receptor, or that multiple defense pathways could be working in concert to provide resistance, similar to our observations with antibacterial defenses under GRP control. Our findings are consistent with the GRP of *P. protegens* protecting against phage infection via coordinated regulation of phage defense systems, akin to its role in defense against interbacterial antagonism.

## Discussion

In this study, we show that the GRP of diverse *Pseudomonas* species coordinates the regulation of pathways for defense against both interbacterial antagonism and phage. The coordinated regulation of antibacterial and antiviral defenses has precedence in more complex organisms; type I interferon signaling in mammals, for example, controls the activation of a wide range of immune responses, including both antiviral and antibacterial mechanisms^69^. The convergent evolution of mechanisms for coordinating responses to disparate threats begs the question as to the adaptive value of such a coupling. Both interferon signaling and the GRP provide a mechanism by which a population of cells can become alerted to the presence of a wide range of threats, before directly encountering the threat itself. In the former case, this is achieved via the numerous receptors for the detection of diverse pathogen-associated molecular patterns that trigger secretion of a common signaling molecule^69^. The GRP, in contrast, is directly activated by the release of cell lysate, a signal that is generated in the course of both interbacterial antagonism and infection by lytic phages^20^. In both situations, by coupling activation of many pathways to a single, diffusible signal, populations gain the ability to anticipate a broad array of attacks and mount a general response, thus sidestepping the limitations of linking defense pathway activation to direct detection of incoming threats within an individual cell.

Our study revealed that the suite of defensive responses within the GRP regulon varies substantially across *Pseudomonas* species. Recognition of the full extent to which these regulons vary was facilitated by our development of a sensitive and comprehensive method for their characterization, which involves comparative proteomic analysis of GRP-activated and inactivated strains. A particular advantage of this approach over some earlier methods is its ability to capture both direct and indirect targets of the pathway^24,32^. The variability we uncovered in the GRP regulon suggests that the specific threats encountered by a given species could be driving the diversification of defensive mechanisms it can produce. This is supported by the observation that different GRP targets in a given species can be important for defense against different antagonists, as observed in studies of *P. aeruginosa^5,26^*.

The widespread conservation of the GRP across the *Pseudomonas* genus has long been at odds with the many conflicting theories put forth regarding its adaptive function in different contexts. In *P. aeruginosa,* for example, the GRP negatively regulates the type III secretion system required for acute infection and positively regulates traits associated with chronic infections, such as extracellular polysaccharide production^25,70^. Accordingly, researchers argue it serves to mediate the switch acute and chronic infection-causing lifestyles^24^. In contrast, in plant protective *Pseudomonas* species, where the pathway is linked to regulation of secondary metabolites important for biocontrol of plant pathogens, it is argued be a form of quorum sensing^71^. Our finding that the GRP coordinates induction of diverse defensive mechanisms in response to antagonism suggests a clear adaptive benefit of the pathway relevant to the physiological conditions shaping the evolution of the genus. Reinterpreting previous work in light of our findings, many prior observations hint at the importance of the GRP in defense against antagonism. For example, early studies on *P. syringae* showed that inactivation of GacS had no impact on its propagation when applied to leaf surfaces under greenhouse conditions, but in field studies, where successful plant colonization requires contending with many competitors, *gacS* mutants were highly impaired^72,73^. Similarly, a previous study found GRP-deficient mutants of *P. protegens* have decreased fitness during interspecies competition, but grow as well as the wild-type in pure culture^74^. Prior studies have also contributed data consistent with the involvement of the GRP in phage defense^75,76^. For example, a group found that in the presence of exogenous putrescine, GRP-regulated polyamine uptake increases resistance of *P. aeruginosa* to several phage^75^. Our results, taken together with these prior findings, strongly support the contention that a key adaptive function of the GRP is to coordinate defensive responses.

The clear benefits conferred by linking defensive mechanisms under a common regulatory pathway raises the question of whether this phenomenon occurs more broadly. While our work suggests that GRP control of defense requires an expanded version of the pathway that is limited to *Pseudomonas* species, the possibility that different accessory kinases could be performing an analogous role in other γ-proteobacteria remains unexplored. Indeed, a recent study employing *Serratia* and *Pectobacterium* strains demonstrated GRP-mediated regulation of CRISPR-Cas systems, suggesting the GRP may have an immune-related role in these organisms^77^. We speculate that pathways coordinating defensive responses in other bacterial phyla may well await discovery.

## Acknowledgements

We thank Donald Kobayashi for kindly providing *Lysobacter enzymogenes* C3, Simon Dove for reagents and helpful discussion, Ricard Rodriguez and Judit Villen for assistance with mass spectrometry, Yaxi Wang and Andi Liu for proteomics experiment guidance, Bob Ernst for assistance with LPS result interpretation, Lydia Contreras and Alex Lukasiewicz for helpful discussions about GRP regulation, the Meeske laboratory for phage experiment guidance and reagents, and members of the Mougous laboratory for insightful discussions. This work was supported by the NIH (5R01AI080609 to J.D.M. and CMB Training Grant T32 GM136534 to D.M.B). J.D.M. is an HHMI investigator and is supported by the Lynn M. and Michael D. Garvey Endowed Chair at the University of Washington.

## Author contributions

D.M.B., S.B.P. and J.D.M. conceived the study. D.M.B., S.K.B., L.A.G., D.Z. and J.D.M. designed the study. D.M.B., S.K.B., L.A.G., A.B., and M.M.d.S. performed experiments. D.M.B., S.K.B., L.A.G., Y.T., D.Z., S.B.P. and J.D.M. processed, analyzed and visualized the data. D.M.B., S.B.P. and J.D.M. wrote the manuscript. D.M.B, S.B.P. and J.D.M. provided supervision and J.D.M. provided funding. All authors contributed to manuscript editing and support the conclusions.

---

**Figure S1.**
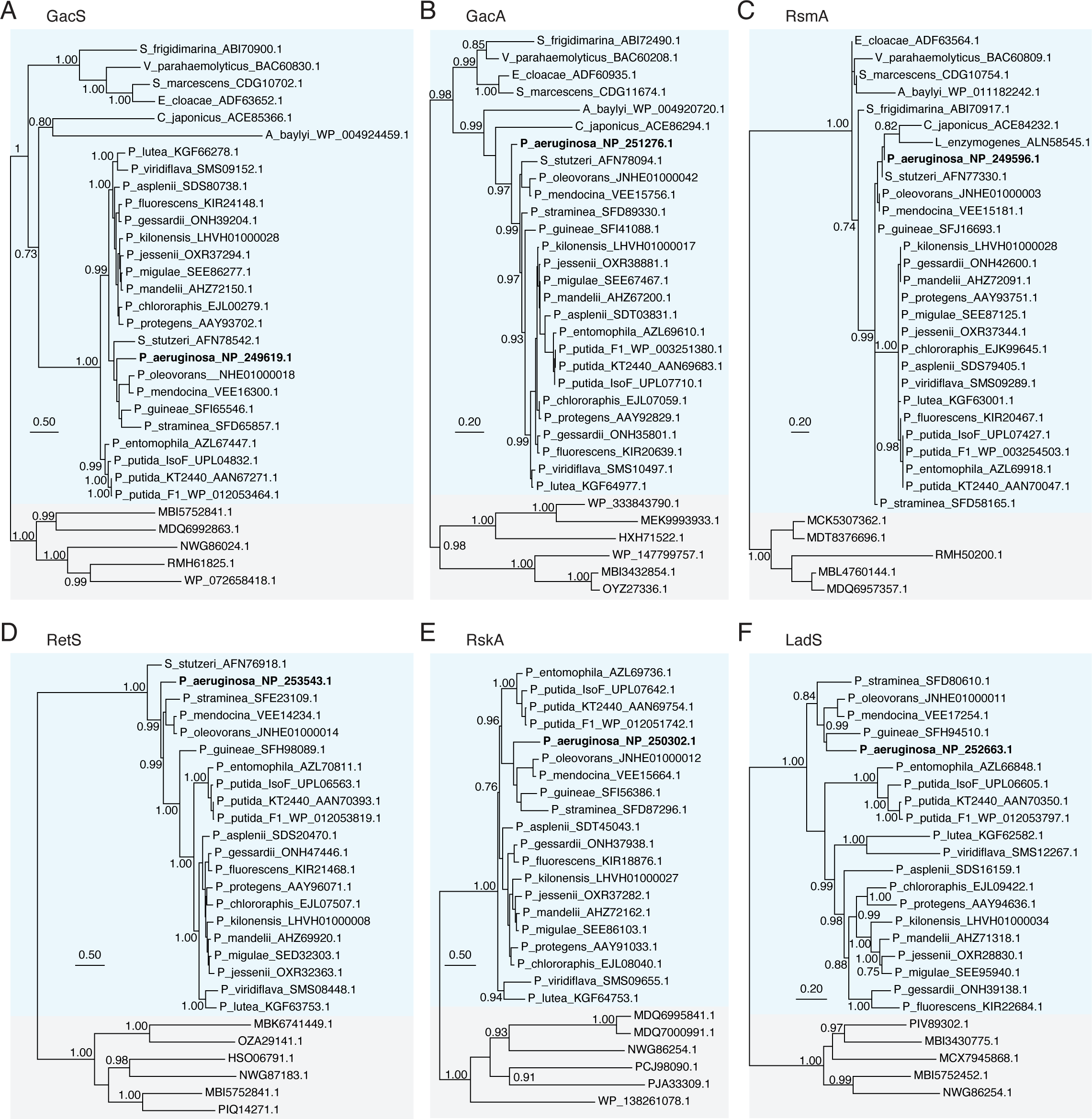
Phylogenetic analysis enables confident identification of GRP signaling protein homologs. (A-F) Maximum-likelihood inferred phylogenies of homologs of the indicated GRP proteins from *P. aeruginosa.* Blue shading indicates orthologous proteins; more distant homologs used as outgroups indicated in grey.

**Figure S2.**
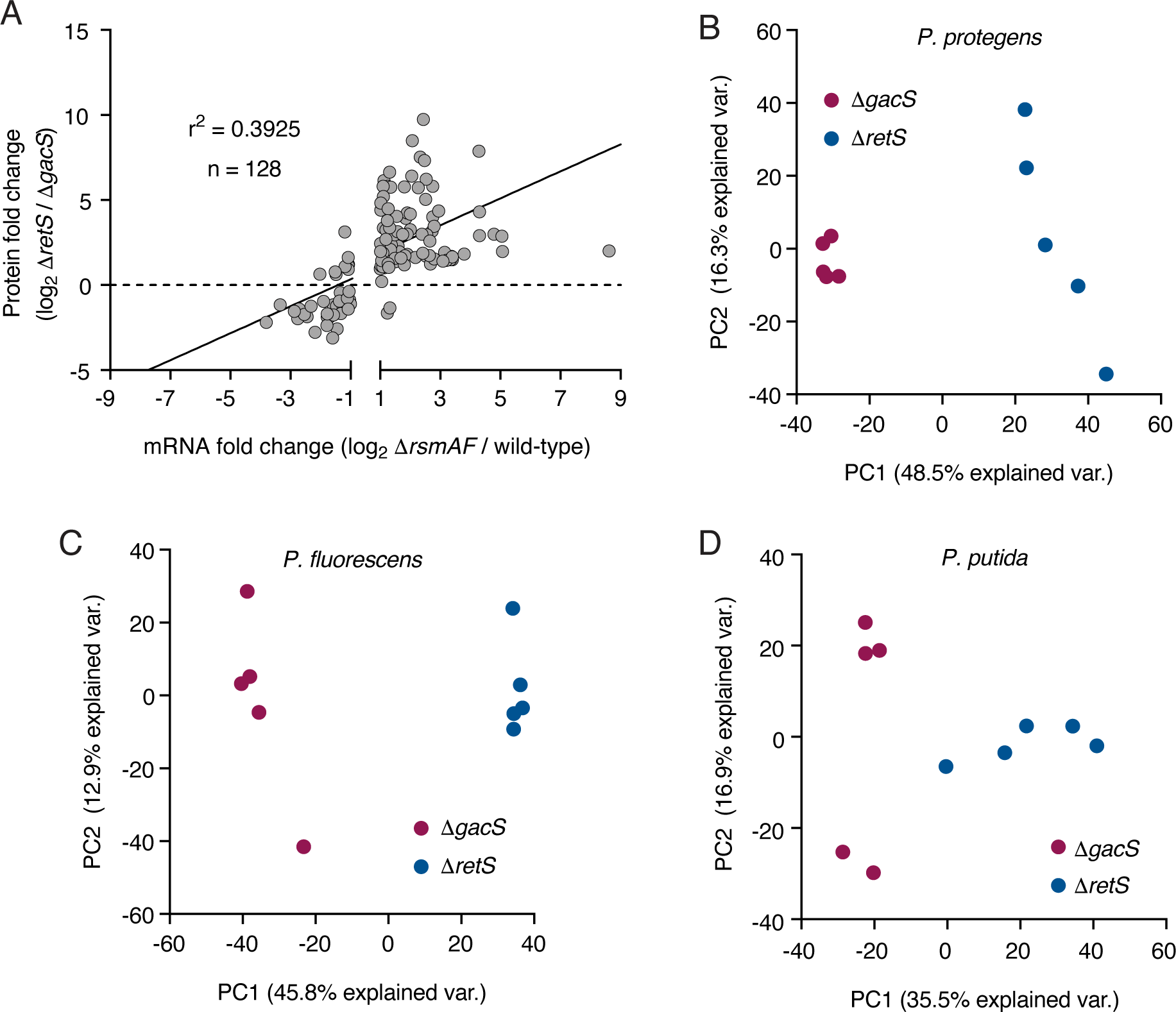
GRP activation yields a replicable shift in the proteomes of *Pseudomonas* species. (A) Comparison of the fold change of protein between *P. aeruginosa* Δ*retS* and Δ*gacS* (this study) with transcript fold change between *P. aeruginosa* Δ*rsmAF* and wild-type from a published RNA-seq dataset^32^. (B-D) Principal component analysis of MS-derived whole-cell proteomes (n=5) from the indicated strains of *P. protegens* (B), *P. fluorescens* (C), and *P. putida* (D).

**Figure S3.**
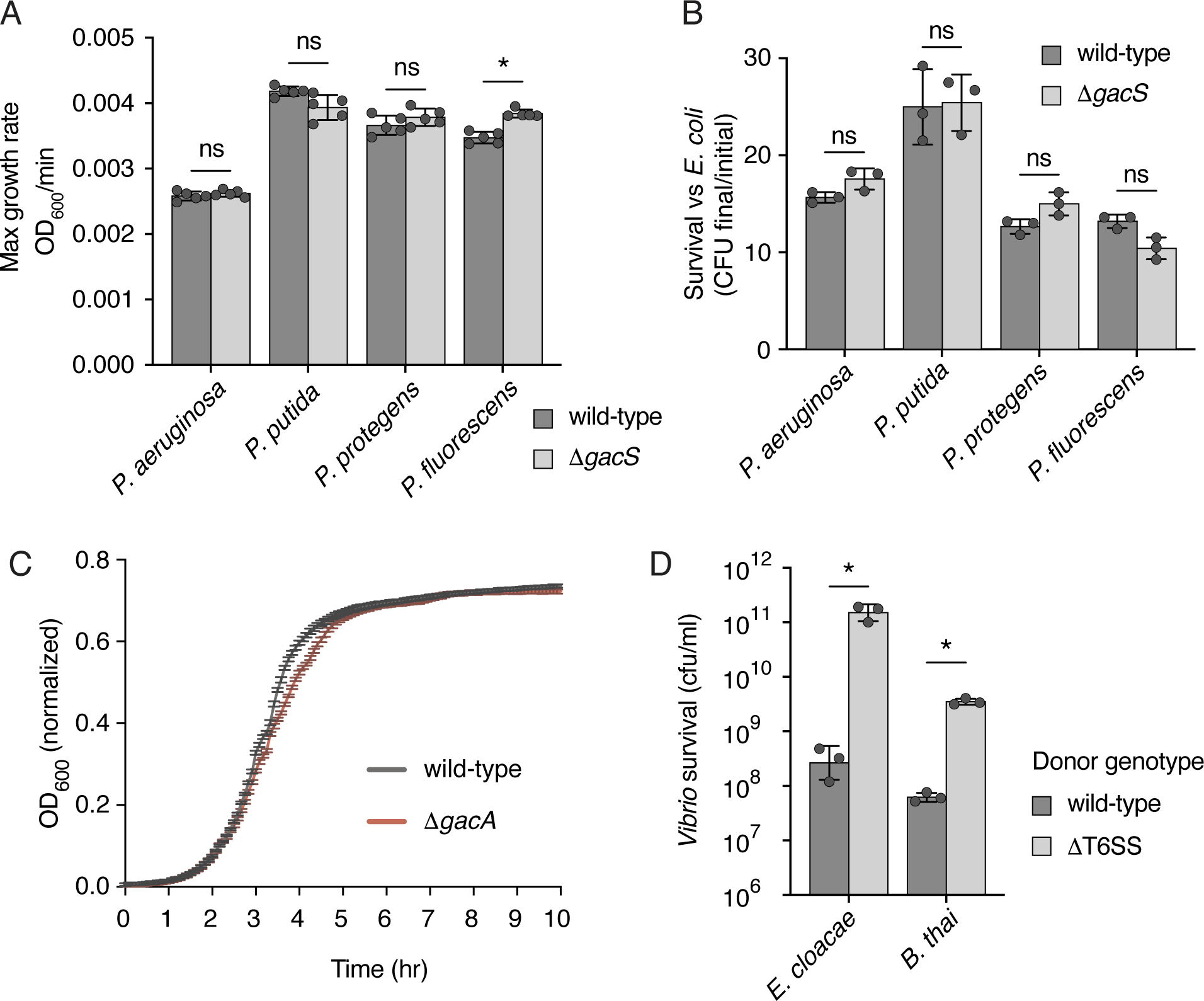
The GRP is dispensable in the absence of antagonism. (A) Comparison of maximum growth rate, defined as the largest change in OD_600_ in a 5-minute interval, between wild-type and *gacS* mutant strains of the indicated *Pseudomonas* species (n=5). (B) Survival of wild-type and *gacS* mutant strains of the indicated *Pseudomonas* species in competition with the non-antagonistic *E. coli* strain MG1655 (n=3). (C) Liquid growth curve measuring OD_600_ of wild-type *V. parahaemolyticus* and a *gacA* mutant strain over time (n=10). OD_600_ is normalized to a blank LB well, and error bars indicate SD. (D) Survival of *V. parahaemolyticus* in competition with the indicated strains of *E. cloacae* and *B. thai*, determined by recovered cfu concentrations (n=3). Statistical significance for all panels was determined using a Welch’s t-test with BH correction for multiple comparisons.

**Figure S4.**
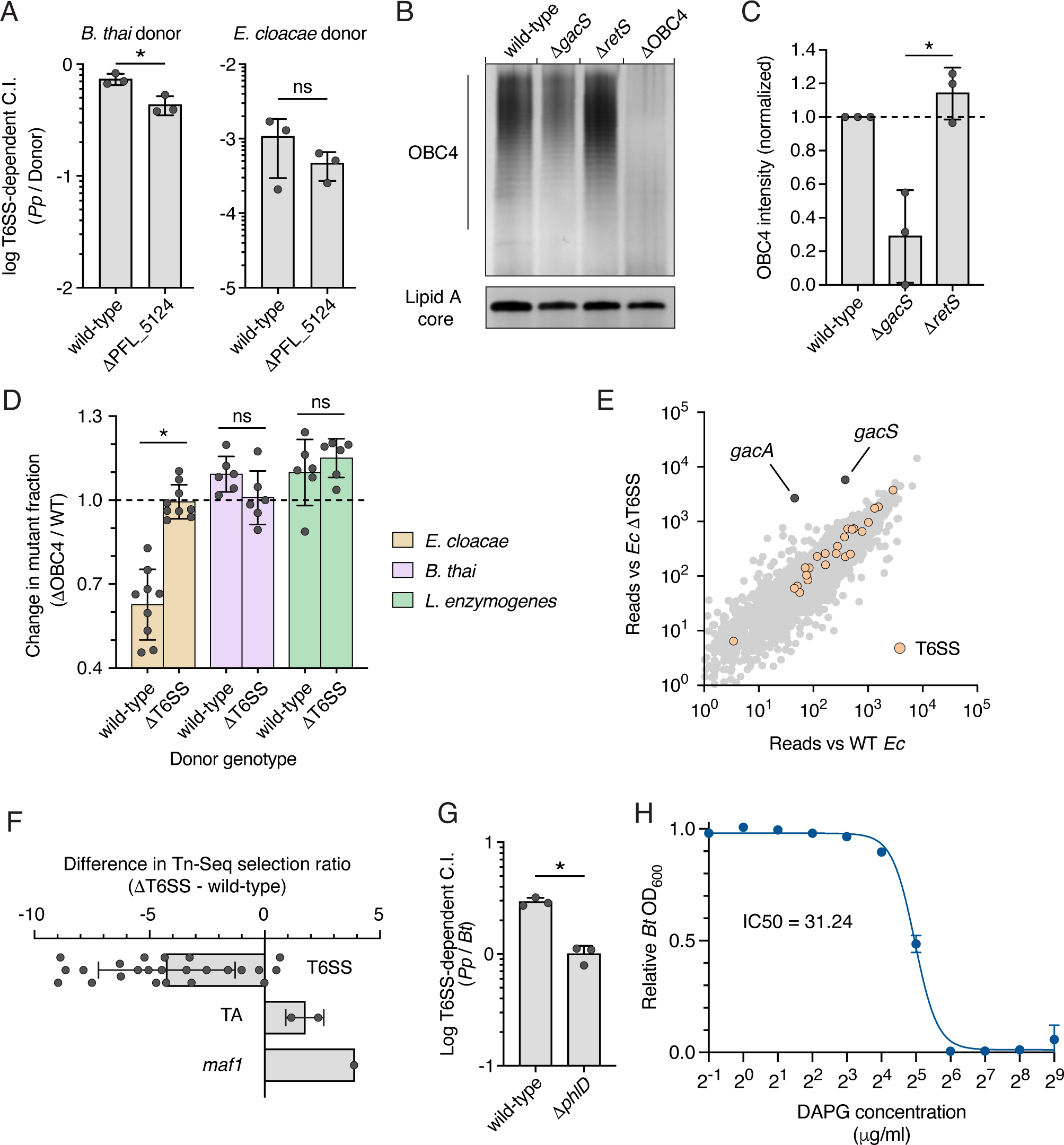
GRP-regulated factors protect against antagonism and their effect can be masked by the T6SS. (A) Competitive fitness of *P. protegens* lacking periplasmic protease PFL_5124 in response to T6SS antagonism by *B. thai* or *E. cloacae* (n=3). (B) Representative LPS profiles of the indicated *P. protegens* strain, normalized by OD_600_, run on an 8-16% gradient polyacrylamide gel and visualized by silver stain. Due to the sensitivity of silver staining, OBC4 and Lipid A core regions shown are taken from separate biological replicates. (C) Densitometry analysis of OBC4 O-polysaccharide from silver-stained gels, normalized to wild-type levels (n=3). (D) Relative survival of *P. protegens* ΔOBC4 in a mixed-recipient competition with the wild-type strain and the indicated strains of *E. cloacae*, *B. thai*, or *L. enzymogenes*. Mutant fraction of the *P. protegens* population normalized to starting conditions, as determined by qPCR (n=6-9). Statistical significance was determined using a Welch’s t-test with BH correction for multiple comparisons. (E) Log-log plot of transposon insertions reads from a *P. protegens* ΔT6SS transposon library competed against wild-type *E. cloacae* (x-axis) or a T6SS deletion strain. (F) Difference in Tn-seq selection ratio (reads vs *Ec* ΔT6SS / reads vs wild-type) for the indicated pathways between the T6SS-inactivated and wild-type transposon libraries. (G) Competitive fitness of the *P. protegens* lacking the capability to produce 2,4-DAPG in response to T6SS antagonism by *B. thai* (n=3). (H) Growth inhibition of *B. thai* by varying concentrations of 2,4-DAPG, as measured by OD_600_ relative to a DMSO control in late log-phase cultures (n=3). Statistical significance for panels A, C, and G was determined using a Welch’s t-test. Asterisks indicate p < 0.05.

**Figure S5.**
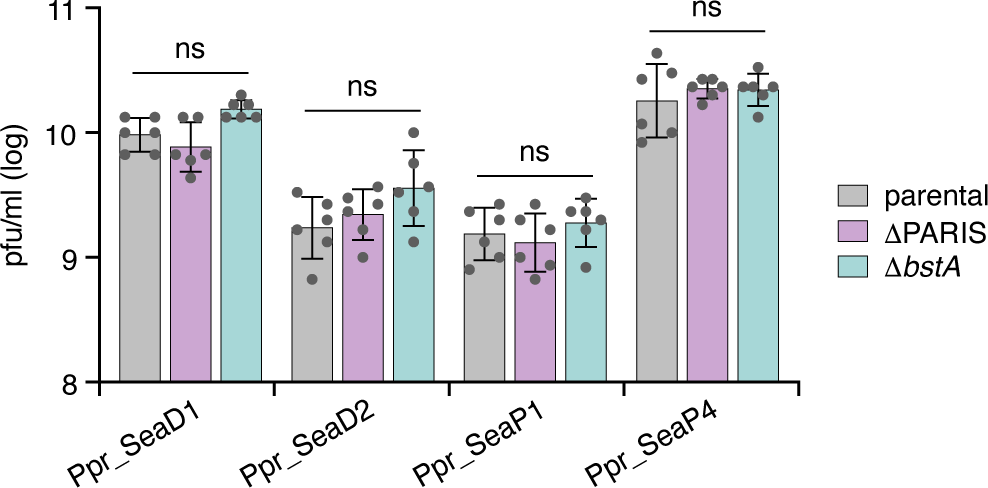
Many *P. protegens* phages are not restricted by PARIS or BstA. Quantification of plaquing efficiency of the indicated phages on mutants of *P. protegens* in a GRP activated (Δ*retS*) background. Statistical significance was assessed for each phage by a one-way ANOVA comparing PFU yield between strains.

## Methods

### Bacterial strains and culture conditions

All strains used in this study are described in Supplementary Table 4 and are available upon request. Strains were derived from *P. aeruginosa* PAO1^78^, *P. putida* IsoF^79^, *P. putida* KT2440^80^, *P. protegens* Pf-5^81^, *P. fluorescens* 2-79^82^, *E. cloacae* ATCC 13047^83^, *B. thai* E264^84^, *L. enzymogenes* C3-1^65^, *V. parahaemolyticus* RIMD 2210633^85^, and *E. coli* MG1655^86^. *E. coli* strains DH5α (Thermo Fisher Scientific) and EC100D pir+ (Lucigen) were used for cloning and maintenance of plasmids. *E. coli* strains S17-1 λpir^87^, Sm10 λpir (Biomedal Lifescience, Cat# BS-3303), HB101^88^, and RHO3^89^ were used for plasmid transfer. Routine growth was performed in Tryptic Soy Broth (TSB, *L. enzymogenes*), marine LB (LB containing 3% w/v NaCl, *V. parahaemolyticus*), Minimal Marine Medium (2% NaCl, 0.4% galactose, 5 mM MgSO_4_, 7 mM K_2_SO_4_, 77 mM K_2_HPO_4_, 35 mM KH_2_PO_4_, and 2 mM NH_4_Cl, *V. parahaemolyticus*), or Lysogeny broth (LB, all other strains), aerated, or on TSB, marine LB, minimal marine medium, or LB plates containing 1.5% w/v agar. *E. coli*, *P. aeruginosa*, *E. cloacae*, and *B. thai* were grown at 37°C, while all other species were maintained at 30°C. Media were supplemented as needed with antibiotics at the following concentrations: gentamicin (15 µg ml^-1^, *E. coli* and *B. thai*, 30 µg ml^-^ _1_, *Pseudomonas* and *V. parahaemolyticus*, 150 µg ml^-1^, *L. enzymogenes*), irgasan (25 µg ml^-1^, *Pseudomonas*), trimethoprim (200 µg ml^-1^, *Pseudomonas*), streptomycin (50 µg ml^-1^, *E. cloacae* and *L. enzymogenes*), kanamycin (50 µg ml^-1^, *L. enzymogenes*), chloramphenicol (25 µg ml^-1^, *E. coli* and *V. parahaemolyticus*), and carbenicillin (150 µg ml^-1^, *E. coli*).

### Plasmid and strain construction

All plasmids used in this study are described in Supplementary Table 4 and are available upon request. Plasmids were generated using the following vector backbones: pRE112 (allelic exchange, *V. parahaemolyticus*), pEXG2 (allelic exchange, all other species), pUC18-mini- Tn7T-Gm-AraE-AraC-pBad (insertion at attTn7 site for complementation, *P. protegens*), and pBT20 (transposon library generation, *P. protegens* and *V. parahaemolyticus*^90-93^. Constructs were designed using Geneious Prime software (v2025.0.3) and primers were synthesized by Integrated DNA Technologies. Plasmids were generated by Gibson assembly^94^ (pUC18T-mini- Tn7T-Gm-AraE-AraC-pBad and pEXG2), *in vivo* homology cloning (pEXG2^95^), or restriction cloning (pRE112). For deletion constructs, 500- or 1000-nucleotide sequences flanking the deleted gene, retaining 10-17 amino acids at both the N- and C-termini, were cloned into vector linearized with SacI-HF and XbaI (pRE112, pEXG2, New England Biosciences) or HindIII-HF and KpnI-HF (pUC18T-mini-Tn7T-Gm-AraE-AraC-pBad, New England Biosciences). Fragments were amplified by PCR from bacterial genomes or synthesized by Twist Biosciences. Complementation plasmids were designed with the coding sequence following the ribosome-binding site from *P. aeruginosa hcp1* cloned into the multiple cloning site of pUC18T-mini- Tn7T-Gm-AraE-AraC-pBad^91^. Assembled plasmids were transformed into CaCl_2_ competent DH5α (pEXG2) or EC100D pir+ (pRE112) *E. coli* by 50 second heat shock at 42°C, recovered for 1 hour in 2XYT, and plated on selective media. Plasmid identity was confirmed by Sanger sequencing (Azenta Life Sciences).

Strains bearing in-frame deletions were generated by allelic exchange, as previously described^93^. Briefly, plasmids were transformed into *E. coli* strains S17-1 λpir (delivery to *L. enzymogenes*) or Sm10 λpir (all other species), then delivered to recipient strains by conjugation. Plate-grown donor and recipient cells were mixed on LB agar, incubated 6 hours at 37°C (*P. aeruginosa*) or 30°C (all other species), and plated on selective media. Transconjugant colonies were counterselected on LB containing 15 (*V. parahaemolyticus*) or 10% sucrose (all other species) at 30°C and deletions were confirmed by sequencing across the insertion junction.

Complementation plasmids (pUC18-mini-Tn7T-Gm-AraE-AraC-pBad) were delivered to recipient cells by tri-parental mating, as previously described^88,96^. Briefly, 100 µL each overnight cultures of Sm10 *E. coli* with the complementation plasmid, recipient strain, *E. coli* HB101 with pRK2013, and *E. coli* Sm10 with pTNS3 were combined, pelleted by centrifugation at 7,000 x g, washed once with fresh LB, and resuspended in 30 µL LB. The mixture was spotted onto LB agar and incubated for 6-8 hours at 30°C. Following incubation, conjugation spots were harvested, resuspended in LB, and plated on selective media. Transconjugant colonies were streaked for isolation a second time, and individual colonies were screened by PCR for the desired insertion.

### Bacterial growth assays

Cultures were grown in liquid LB from isolated colonies, with shaking, at 37°C (*P. aeruginosa*) or 30°C (all other strains) until turbid. Cultures were then diluted to OD_600_ = 0.01, transferred to a CellStar clear bottom 96-well plate (Greinier), with 5-10 replicates per genotype, and sealed with a BreatheEasy membrane (Diversified Biotech). Plates were incubated in a LogPhase 600 Microbiology Reader (BioTek) set to 37°C (*P. aeruginosa*) or 30°C (all other strains), shaking at 800 rpm, with OD_600_ measurements taken every 5 minutes for a total of 18 hours. Maximum growth rate was determined by the LogPhase software using the change in OD_600_ reading in successive measurements during log phase growth. Maximum growth rates were compared using a Welch’s t-test with Benjamini-Hochberg (BH) correction for multiple comparisons.

### Competitive growth assays

Strains were grown overnight in liquid media, with shaking, inoculated from single colonies from freshly streaked plates. Cultures were collected with recipient strains (*Pseudomonas*, *V. parahaemolyticus*) in mid-log phase growth and donor strains (*E. cloacae*, *L. enzymogenes*, *B. thai*, and *E. coli*) in early stationary phase, pelleted by centrifugation at 4,000 x g for 10 minutes, resuspended at high cell density (OD_600_ = 100) in fresh LB media, and mixed together at defined OD ratios. For *gacS* and *gacA* mutant competitions, cultures were mixed with *E. cloacae* at a donor:recipient OD ratio of 8:1 (*P. aeruginosa* and *V. parahaemolyticus*) or 4:1 (*P. putida*, *P. putida,* and *P. fluorescens*), or were mixed with *B. thai* at a donor:recipient ratio of 1:4 (*P. fluorescens*), 50:1 (*V. parahaemolyticus*), or 4:1 (all other strains). For the PFL_5124 mutant competition, cultures were mixed at a donor:recipient ratio of 10:1 (*E. cloacae*) or 20:1 (*B. thai*). 5 µl spots were plated on LB with 3% (w/v) agar (*V. parahaemolyticus*) or no-salt LB with 3% (w/v) agar (all other recipients) and were incubated at 6 hours at 37°C (*P. aeruginosa*) or 30°C (all other recipients). Surviving CFUs were enumerated by serial dilution on selective media. Contribution of genes to antagonism-dependent fitness was determined by the relative antagonism-dependent competitive index ([CI_WT_vs_WT_donor_ / CI_WT_vs_ΔT6SS_donor_] / [CI_mutant_vs_WT_donor_ / CI_mutant_vs_ΔT6SS_donor_]) of each strain. Competitive index is defined by [recipient_final_ / donor_final_] / [recipient_initial_ / donor_initial_] using recovered CFU counts. Data were analyzed using R v4.4.0 ^97^ and plotted using Prism (GraphPad, v10.4.1). Statistical significance was determined by Welch’s t-test comparing log transformed T6SS-dependent competitive index between wild-type and mutant recipients, with BH correction for multiple comparisons.

Competitive growth assays with OBC4 mutant strains were performed as described above, with the following modifications. *P. protegens* recipient cells were resuspended directly from LB agar plates into 10 mL LB. Donor and recipient cultures were normalized to OD_600_ = 1 and 0.1, respectively. Following competition, genomic DNA of recovered cells was purified using a QIAGEN Blood and Tissue gDNA prep kit and relative proportions of wild-type *P. protegens* and *P. protegens* ΔOBC4 were determined by qPCR on a CFX Connect Real-Time System (Bio-Rad). Significant differences in mutant fraction change were determined by Welch’s t-test, with BH correction for multiple comparisons.

### Proteomics

*Pseudomonas* cultures were grown in 2 ml liquid LB media, with shaking, at 30°C (*P. protegens* and *P. fluorescens*) or 37°C (*P. aeruginosa*) until late log phase growth, determined by an OD_600_ between 1.2 and 1.5. Cells were pelleted by centrifugation at 9,000 x g for 3 minutes, washed twice in phosphate buffered saline (PBS), resuspended in 100 µl lysis buffer (8 M urea, 75 mM NaCl, 50 mM Tris-HCl, pH 8.2), lysed by three freeze-thaw cycles interspersed with sonication in a Cole-Parmer ultrasonic cleaner, and centrifuged at 20,000 x g for 10 minutes. Protein content in the supernatant was quantified using a Pierce BCA Protein Assay Kit (Thermo Fisher Scientific). Samples were reduced in 5 mM 1,4-dithioreitol (DTT) for 25 minutes at 55°C, cooled to room temperature, and alkylated using 14 mM iodoacetamide for 30 minutes in the dark. Alkylation was quenched by addition of DTT to 10 mM. Samples were diluted in 5 volumes of 25 mM Tris-HCl, pH 8.2, CaCl_2_ was added to 1 mM, and protein was digested at 37°C overnight with 4 µg/ml trypsin. Digestion was halted with trifluoroacetic acid (TFA, 0.4% final volume), samples were centrifuged 10 minutes at 20,000 x g, and pellets were discarded, retaining supernatant. *P. aeruginosa* and *P. protegens* samples were loaded onto BioPureSPN Mini RPC desalting columns (Nest Group) preconditioned with 100% acetonitrile (ACN), LC- MC grade water, and 0.1% formic acid (FA), washed with 0.1% FA, and eluted in 80% ACN with 0.1% FA. For *P. fluorescens*, 0.4% TFA was added to samples, which were then applied to MCX StageTips^98^ that had been preconditioned twice with 100% MeOH, once each with 100% ACN, 75% ACN with 5% NH_4_OH, and 75% ACN with 0.5% acetic acid, and twice with 0.1% TFA. StageTips were washed with 0.1% TFA, twice with 75% ACN containing 0.5% acetic acid, and once with 0.5% acetic acid in LC-MS grade water. Samples were eluted in 75% ACN with 5% NH_4_OH. All samples were dried using an SPD10130 SpeedVac (Thermo Fisher Scientific) and resuspended in 5% acetonitrile with 0.1% formic acid (FA) to a final protein concentration of 0.25 µg/ml. Samples were analyzed by LC-MS/MS as previously described^5^ using a Lumos Fusion Orbitrap Mass Spectrometer (Thermo Fisher Scientific). Spectra were mapped to reference proteomes and analyzed using the MaxLFQ algorithm in MaxQuant software^99,100^, using 1% FDR cutoffs at the peptide, protein, and site levels. The unannotated ORF downstream of PFL_2023 predicted as phage defense gene *ariB* was not included in the initial peptide matching. Peptide abundance for this protein was determined by a second MaxQuant run. Missing values were imputed from a normal distribution of log-transformed data defined by a mean at (data median – 1.4 x std. dev) and a standard deviation of (data std. dev * 0.5), akin to previously described methods^101^. Differentially regulated proteins were identified using a Welch’s t-test with BH correction for multiple comparisons and a filter of >2-fold difference between genotypes. Principal component analysis was performed using the ggbiplot package in R ^102^. Identification of phage defense systems was performed using web tools PADLOC and DefenseFinder^103,104^.

### Transposon library generation

A Himar1 transposon insertion library was generated in *P. protegens* Pf-5, Pf-5 Δ*tssM*, and *V. parahaemolyticus* RIMD 2210633 using established methods^5^. Briefly, the suicide vector pBT20, containing a Himar1 mariner transposon and transposase, was conjugated into recipient strains from *E. coli* Sm10 λpir for 6 hours on LB agar (*P. protegens*) or from RHO3 overnight on LB agar containing 400 µg/ml diaminopimelic acid (DAP) (*V. parahaemolyticus*). Transconjugant colonies were selected on LB agar containing 30 µg/ml gentamicin and 25 µg/ml irgasan (*P. protegens*) or on marine LB agar containing 30 µg/ml gentamicin (*V. parahaemolyticus*), then pooled in LB containing 15% dimethylsulfoxide (DMSO, *P. protegens*) or 15% glycerol (*V. parahaemolyticus*). Libraries were aliquoted, flash frozen in liquid nitrogen, and stored at -80°C. Transposon insertion sequencing, as described below, revealed library complexities of 148,362 unique insertion sites in the wild-type *P. protegens* library, 170,203 in the *P. protegens* Δ*tssM* library, and 304,000 in the *V. parahaemolyticus* library.

### Tn-seq screen and analysis

Himar1 mariner transposon mutant libraries were grown to log phase in LB broth, shaking, at 30°C. Cells were pelleted by centrifugation at 4,000 x g for 10 minutes, washed in fresh LB and resuspended at a defined OD_600_ to yield the donor:recipient ratio described below when mixed with a donor strain at OD_600_ = 100. 196 replicate spots of 5 µl of each mixture were plated on marine LB with 3% (w/v) agar (*V. parahaemolyticus*) or on no-salt LB with 3% (w/v) agar (*P. protegens*) and incubated at 30°C for a defined time period described below. The wild- type *P. protegens* library was grown with *E. cloacae* (wild-type or Δ*tssM*) at a 100:12.5 donor:recipient ratio for 18 hours, *B. thai* (wild-type or Δ*tssM*) at 100:2 for 18 hours, and *L. enzymogenes* C3 (wild-type or Δ*virD4*) at 100:12.5 for 6 hours. The *P. protegens* Δ*tssM* library was grown with *E. cloacae* (WT or Δ*tssM*) at a 100:25 donor:recipient ratio for 18 hours. The *V. parahaemolyticus* library was grown with *E. cloacae* (WT or Δ*tssM*) at a 100:12.5 donor:recipient ratio for 4 hours. Following incubation, cell mixtures were resuspended in LB and immediately plated for surviving *Pseudomonas* and *Vibrio* cells on LB agar containing 25 µg/ml irgasan (*P. protegens*) or on LB agar containing 30 µg/ml gentamicin (*V. parahaemolyticus*). After overnight incubation at 30°C (*P. protegens*) or 20°C (*V. parahaemolyticus*), cells were harvested using a sterile cell scraper from plates flooded twice with 10 mL LB, and genomic DNA was purified using the QIAGEN Blood & Tissue gDNA prep kit. Tn-seq libraries were generated by DNA shearing, C-tailing, and repeated PCR amplification, as previously described^105^. Transposon insertion libraries for each experiment were pooled, and multiplexed samples were sequenced as 51-base single-end reads on an Illumina MiniSeq with 30-40% phiX DNA spiked in.

Illumina sequencing reads were analyzed using a previously described custom Python script^106^. Briefly, sequences were filtered for those with the first 6 bases matching the transposon end and were mapped to the relevant genome using a BWA aligner. Reads per insertion site were tallied, and the sum of all insertion sites within 5 and 90% of a coding sequence was recorded for each gene. Transposon insertion site reads for a library competed against different donor strains were normalized by the median fold-difference between all sequenced genes. Missing values were imputed from a normal distribution of log-transformed data defined by a mean at (data median – 1.4 x std. dev) and a standard deviation of (data std. dev * 0.5). Conditional gene essentiality was determined using a Mann-Whitney U-test comparing the fold-difference in reads between donor strains for each transposon insertion site in a gene with the same ratio across the whole genome, using BH correction for multiple comparisons.

### Identification and phylogenetic analysis of Gac/Rsm system orthologs

Orthologous gene groups (orthogroups) were identified by integration of two complementary methods. First, orthologous relationships between proteins were identified across diverse bacterial genomes using OrthoFinder v2.2.3^107^. In parallel, an independent all-against-all BLASTP search^108^, with a significance threshold of 0.005, was performed. Only reciprocal top hits from different genomes were grouped into orthogroups. Results from both methods were integrated, and false positives were further removed by domain architecture analysis, gene neighborhood analysis, and phylogenetic inference. For domain architecture analysis, hmmscan^109^ was used to search both the Pfam database^110^ and a custom in-house domain profile database. Gene neighborhood was assessed by examining the ten genes upstream and downstream of each target gene. Phylogenetic relationships were inferred using PhyML^111^.

### Generation of GTDB-based phylogeny and cladogram

The phylogenetic tree underlying the Genome Taxonomy Database (GTDB) v220 was downloaded from the GTDB^112-115^ website (https://gtdb.ecogenomic.org/) and converted to Newick format in Geneious Prime (v2025.0.3) Select reference species were pruned from the tree using the prune function in ETE software v3.1.3^116^, preserving branch length. Trees were visualized and converted to cladograms, where appropriate, using FigTree (v1.4.4). Species were labeled using NCBI taxonomy.

### LPS extraction and quantification

LPS extraction and visualization was performed using established methods^117,118^. Briefly, overnight cultures were pelleted by centrifugation at 9,000 x g for 3 minutes, washed twice in PBS, and resuspended at a concentration of OD_600_ = 0.75. Serial dilutions were plated on LB agar to ensure equal cell concentrations. 1 mL of resuspended cells was centrifuged 10 minutes at 10,600 x g, discarding supernatant. Pellets were optionally stored at -20°C. Cells were gently resuspended in 200 µl SDS buffer, freshly diluted in distilled H_2_O from a 2x stock (4% β-mercaptoethanol, 4% SDS, 20% glycerol in 0.1 M Tris-HCl, pH 6.8) and boiled at 95°C for 10 minutes. Samples were cooled to room temperature for 10 minutes, then 5 µl of RNase A (10 mg/ml) and 2 µl of Turbo DNase (Invitrogen) were added. Samples were gently mixed and were incubated at 37°C for 30 minutes. Following nuclease treatment, 10 µl of proteinase K (10 mg/ml) was added, and samples were incubated at 59°C for 3-4 hours, shaking at 350 rpm. 10 µl of each sample was loaded onto an 8-16% gradient Criterion TGX stain-free precast polyacrylamide gel (Bio-Rad), and a 100V current was applied until the dye front reached nearly to the bottom of the gel. Gels were briefly rinsed in dH_2_O, incubated 15 minutes on an orbital shaker with fixing-oxidizing solution (40% ethanol, 5% acetic acid, 1% sodium (meta)periodate), then silver stained with a SilverQuest staining kit (Invitrogen) according to the manufacturer’s protocol, beginning with the sensitization step. Gels were imaged using a GelDoc Go imager (Bio-Rad). Due to the sensitivity of silver staining and the differential abundance of OBC4 O- polysaccharide and Lipid-A core, the two gels shown in Supplementary Figure 4B derive from distinct biological replicates.

OBC4 was quantified by densitometry using FIJI^119^, according to established methods^120,121^. Briefly, a consistently sized box was drawn to one-third the width of each lane at the OBC4 region of the gel and mean signal intensity inside the box was measured. The lane containing *P. protegens* ΔOBC4 was set as background. Mean signal intensity of background was subtracted, and samples were normalized to the wild-type lane. Statistical significance was determined by a Welch’s t-test.

### 2,4-DAPG sensitivity assay

Bacterial sensitivity to 2,4-DAPG was determined using established methods^122^. Briefly, cultures were grown to early log phase from freshly streaked colonies, diluted to OD_600_ = 0.001, and pipetted into a clear-bottom 96-well plate (150 µl/well). 4 µl of 2,4-DAPG, serially diluted two-fold in DMSO, was added to each well to the indicated final concentration. Cultures were grown in triplicate for 24 hours in a LogPhase 600 Microbiology Reader (BioTek) set to 30°C, shaking at 800 rpm, with OD_600_ measurements collected at 5 minute intervals. OD_600_ readings were corrected by subtraction of background from a blank LB well. Sensitivity was determined by the OD_600_ reading relative to a DMSO control as the DMSO control was exiting log phase growth (10.08 hours). IC50 was calculated based on a sigmoidal fit of the relative OD_600_ readings.

### Phage isolation and propagation

Phages were isolated from fresh rhizosphere soil samples collected from Denny Park or the Picardo Farm P-Patch Community Garden, both located in Seattle, Washington. Soil, including pieces of plant roots, was resuspended in 15 mL PBS and shaken at room temperature for one hour on an orbital shaker (Thermolyne). The resulting suspension was centrifuged 10 minutes at 2,000 x g, and supernatant was filtered sequentially through a 100 µm cell strainer (Corning) and a 0.45 µl syringe filter (Thermo Scientific). 300 µl filtrate was incubated 10 minutes at room temperature with 50 µl of a log phase culture of *P. protegens* Δ*gacS*, supplemented with 5 mM CaCl_2_, mixed with 3 ml LB top agar containing 0.5% agar, poured evenly onto LB agar plates, and incubated overnight at 30°C. Plaques were picked with a 20 µl barrier pipet tip, resuspended in 500 µl SM buffer (50 mM Tris-HCl, pH 7.5, 100 mM NaCl, 8 mM MgSO_4_), and filtered through a 0.22 µm syringe filter (Thermo Scientific). Each phage was plaque purified in this manner at least twice to ensure stock purity. To generate phage stocks, 5 ml SM buffer was added to top agar following complete or near complete bacterial cell clearance. Plates were sealed with parafilm and rocked overnight at 4°C to release phage particles. Buffer was collected from the flooded plate, centrifuged 8 min at 3,000 x g, and supernatant was serially filtered through 0.45 µm and 0.22 µm syringe filters. Stocks were stored at 4°C.

Plaquing efficiency for each phage was determined by spotting 10-fold serial dilutions of a single phage stock onto top agar lawns containing different *P. protegens* mutants. *P. protegens* strains were grown overnight on LB agar, resuspended in LB at OD_600_ = 1, mixed with LB top agar containing 5 mM CaCl_2_ (112.5 µl cell suspension in 6.75 ml top agar), and poured evenly onto 15 cm LB agar plates. Top agar lawns were dried 15 minutes in a biosafety cabinet, then 3 µl spots of phage stock, serially diluted in SM buffer, were dispensed using a Rainin BenchSmart pipettor (Mettler Toledo). Plates were incubated at 30°C for 6-15 hours, until plaques appeared. Differential plaquing efficiency of each phage on *P. protegens* Δ*gacS* versus Δ*retS* was determined using a one-sample t-test. Differential plaquing of SeaP2 on phage defense system mutant strains was determined by one-way ANOVA followed by Holm-Sidak post hoc test. Plaque images were captured on a GelDoc Go imager (Bio-Rad). Images were inverted, scaled, and cropped in FIJI.

### Phage sequencing, annotation, and classification

Phage DNA was purified for sequencing from high titer stocks. DNA was released from capsids by sequential incubation with nucleases (100 µg/ml DNase I and RNase I, 37°C for 30- 45 minutes), 10 mM EDTA (37°C for 15 minutes), and 200 µg/ml proteinase K (50°C for 30 minutes). Samples were centrifuged at 21,000 x g for 2 minutes. DNA was precipitated from supernatant by addition of 1/10 volume sodium acetate, pH 5.5, 1/100 volume GlycoBlue, and 2.5x volume 100% ethanol, followed by incubation for 1 hour at -20°C. DNA was pelleted by centrifugation at 21,000 x g for 15 minutes and washed sequentially with 100% isopropanol and 70% ethanol. Pellets were dried 20-30 minutes and gently resuspended in water. DNA concentration and integrity were determined by Qubit fluorometry and gel electrophoresis, respectively.

Sequencing libraries were prepared from 100 - 300 ng purified DNA using Illumina DNA Prep kit. Libraries were pooled and sequenced in multiplex as 2 x 150-bp paired-end reads using an Illumina iSeq. Reads were trimmed using Trimmomatic (Galaxy Version 0.39+galaxy2) with Illuminaclip, Trailing (minimum quality 25-30), AvgQual (minimum 25-30) and MinLen (120 bp) operations^123^. Trimmed paired reads were assembled using SPAdes (Galaxy version 3.15.5+galaxy2) with either automatic or user-specified (55, 77, 99, 101) k-mer size values to obtain a single large contig of >40kb for each purified phage^124^. Phage were taxonomically classified by BLASTP searches of their tail protein sequences against the NCBI nr database. The closest matches corresponded to tail proteins from Pseudomonas phages vB_PpuP-Luke-3 (95.7% aa ID to the B2 tail protein) and PollyC (99.6% aa ID to the Ppr_SeaP2 tail protein). VIRIDIC analysis with the full sequences of these phages indicated that they belonged to the same two genera as those of the phage we isolated (SeaD1-2 with vB_PpuP-Luke-3 and Ppr_SeaP2-5 with PollyC)^125^.

