## Supplementary Tables 3 and 4 for "Pseudomonads coordinate innate defense against viruses and bacteria with a single regulatory system"

**Table S3:** Summary of phage defense systems, related to Figure 2

| Organism | Locus Tag | Defense System | Gene | GRP fold activation | Adj. p-value |
|---|---|---|---|---|---|
| *P. aeruginosa* | PA0574 | SoFic | SoFic | NA | NA |
| | PA0715 | Retron_I-B | RT_I-B | 1.005 | 0.977 |
| | PA0716 | Retron_I-B | ATPase-Toprim_I-B | 0.352 | 0.414 |
| | PA1370 | PDC-S39 | PDC-S39 | 1.433 | 0.423 |
| | PA1371 | Helicase-DUF2290 | DUF2290 | NA | NA |
| | PA1372 | Helicase-DUF2290 | Helicase | 0.896 | 0.185 |
| | PA1671 | PD-T4-6 | PD-T4-6 | NA | NA |
| | PA1935 | Gabija | GajB | NA | NA |
| | PA1939 | Gabija | GajA | 4.940 | 8.09E-03 |
| | PA2732 | RM_type_I | REase_I | 1.11 | 0.325 |
| | PA2734 | RM_type_I | Specificity_I | 0.659 | 0.0432 |
| | PA2735 | RM_type_I | MTase_I | 0.735 | 0.0443 |
| *P. protegens* | PFL_1011 | PDC-S04 | PDC-S04 | 0.802 | NA |
| | PFL_1398 | PDC-S21 | PDC-S21 | 0.914 | 0.320 |
| | PFL_2023 | PARIS_I | AriA | 172.9 | 4.75E-03 |
| | PFL_2561 | PDC-S02 | PDC-S02 | NA | NA |
| | PFL_2676 | PDC-S12 | PDC-S12 | 0.989 | 0.923 |
| | PFL_2962 | Septu_type_I | PtuB1 | NA | NA |
| | PFL_2963 | Septu_type_I | PtuA1 | 1.035 | 0.950 |
| | PFL_2964 | DMS_other | Specificity_I | 1.796 | 0.150 |
| | PFL_2965 | DMS_other | MTase_I | 3.028 | 0.0212 |
| | PFL_3013 | Druantia_type_II | DruM2 | 5.100 | 0.0734 |
| | PFL_3014 | Druantia_type_II | DruF2 | 1.608 | NA |
| | PFL_3015 | Druantia_type_II | DruG2 | 5.454 | 0.0095 |
| | PFL_3016 | Druantia_type_II | DruE2 | 16.903 | 3.87E-06 |
| | PFL_4270 | PDC-S08 | PDC-S08 | NA | NA |
| | PFL_6252 | BstA | BstA | 29.729 | 4.86E-05 |
| *P. fluorescens* | PFLU3_01560 | PDC-S21 | PDC-S21 | 0.449 | 0.111 |
| | PFLU3_04460 | RM_type_IV | mREase_IV | 7.926 | 1.06E-05 |
| | PFLU3_11630 | SoFic | SoFic | 0.687 | 0.315 |
| | PFLU3_12200 | Hachiman_type_I | HamA1 | 6.673 | 1.409E-04 |
| | PFLU3_12210 | Hachiman_type_I | HamB1 | NA | NA |
| | PFLU3_12220 | DRT_class_III | RT_UG5-nitrilase | NA | NA |
| | PFLU3_12490 | PDC-S04 | PDC-S04 | NA | NA |
| | PFLU3_14030 | SoFic | SoFic | NA | NA |
| | PFLU3_18670 | AbiE | AbiEii | NA | NA |
| | PFLU3_18680 | AbiE | AbiEi | NA | NA |
| | PFLU3_21860 | Kiwa | KwaB | 20.798 | 1.478E-04 |
| | PFLU3_21870 | Kiwa | KwaA | NA | NA |
| | PFLU3_21970 | Druantia_type_III | DruE3 | 2.034 | 3.956E-04 |
| | PFLU3_21980 | Druantia_type_III | DruH3 | NA | NA |
| | PFLU3_21990 | RM_type_I | REase_I | 1.354 | 0.405 |

| Organism | Locus Tag | Defense System | Gene | GRP fold activation | Adj. p-value |
|---|---|---|---|---|---|
| *P. fluorescens* | PFLU3_22000 | PDC-S14 | PDC-S14 | 0.402 | 0.0209 |
| | PFLU3_22010 | RM_type_I | Specificity_I | 0.456 | 0.0307 |
| | PFLU3_22030 | RM_type_I | MTase_I | 0.701 | 2.779E-03 |
| | PFLU3_28130 | PDC-S24 | PDC-S24 | 0.606 | 0.250 |
| | PFLU3_30380 | Bunzi | BnzA | 5.815 | 9.754E-04 |
| | PFLU3_30390 | Bunzi | BnzB | 0.967 | 0.967 |
| | PFLU3_31980 | Shango | SngC | NA | NA |
| | PFLU3_31990 | Shango | 3S | 23.817 | 5.151E-04 |
| | PFLU3_32000 | Shango | SngA | NA | NA |
| | PFLU3_37090 | DRT_class_III | Drt1a | NA | NA |
| | PFLU3_37260 | DMS_other | BrxHI | NA | NA |
| | PFLU3_37300 | DMS_other | DrmC | NA | NA |
| | PFLU3_41100 | PD-T7-4 | PD-T7-4 | 7.004 | 2.48E-05 |
| | PFLU3_42830 | PD-T4-6 | PD-T4-6 | NA | NA |
| | PFLU3_45960 | PDC-S08 | PDC-S08 | NA | NA |
| | PFLU3_49280 | SoFic | SoFic | NA | NA |
| | PFLU3_55230 | PDC-S12 | PDC-S12 | 0.762 | 0.185 |
| | PFLU3_56560 | Mokosh_TypeII | MkoC | 9.009 | 1.817E-04 |
| *P. putida* | PP_0049 | PD-T7-1 | PD-T7-1 | NA | NA |
| | PP_1161 | PDC-S12 | PDC-S12 | 0.851 | 0.204 |
| | PP_1406 | PDC-S21 | PDC-S21 | 0.871 | 0.528 |
| | PP_2277 | PDC-S58 | PDC-S58 | NA | NA |
| | PP_2531 | PDC-S08 | PDC-S08 | NA | NA |
| | PP_3680 | Gabija | GajA | 6.994 | 7.915E-04 |
| | PP_3681 | Gabija | GajB | NA | NA |
| | PP_3692 | PDC-S06 | PDC-S06 | NA | NA |
| | PP_3694 | Wadjet_type_I | JetD1 | NA | NA |
| | PP_3695 | Wadjet_type_I | JetD1 | NA | NA |
| | PP_3696 | Wadjet_type_I | JetC1 | NA | NA |
| | PP_3697 | Wadjet_type_I | JetC1 | NA | NA |
| | PP_3698 | Wadjet_type_I | JetA1 | NA | NA |
| | PP_3708 | PDC-S64 | PDC-S64 | NA | NA |
| | PP_3988 | RM_Type_II | Type_II_Rease | 3.883 | 5.233E-03 |
| | PP_3989 | RM_Type_II | Type_II_Mtase | 1.023 | 0.768 |
| | PP_4447 | GAO_20 | DUF4297 | 2.409 | 0.0397 |
| | PP_4448 | GAO_20 | HerA | 8.982 | 2.41E-05 |
| | PP_4740 | RM_type_I | REase_I | 4.544 | 0.0121 |
| | PP_4741 | RM_type_I | MTase_I | 0.994 | 0.949 |
| | PP_4742 | RM_type_I | Specificity_I | 0.995 | 0.973 |
| | PP_5622 | Wadjet_type_I | JetB1 | NA | NA |
| | PP_5643 | PDC-M53 | PDC-M53A | 2.768 | 1.721E-04 |
| | PP_5644 | PDC-M53 | PDC-M53B | NA | NA |

**Table S4:** Strains and plasmids used in this study

| **Strains** | | |
|---|---|---|
| Organism | Genotype | Source |
| *P. aeruginosa* PAO1 | parental | 1 |
| | Δ*gacS* (PA0928) | 2 |
| | Δ*retS* (PA4856) | 3 |
| *P. protegens* Pf-5 | parental | 4 |
| | Δ*gacS* (PFL_4451) | This study |
| | Δ*retS* (PFL_0664) | This study |
| | ΔPFL_5124 | This study |
| | ΔPARIS (PFL_2023 and downstream ORF) | This study |
| | Δ*bstA* (PFL_6252) | This study |
| | ΔPARIS attTn7::AraE-AraC-pBad-PARIS | This study |
| | Δ*bstA* attTn7::AraE-AraC-pBad-*bstA* | This study |
| | Δ*phlD* (PFL_5957) | This study |
| | ΔOBC4 (PFL_5483 - PFL_5495) | This study |
| *P. fluorescens* 2-79 | parental | 5 |
| | Δ*gacS* (PFLU3_03780) | This study |
| | Δ*retS* (PFLU3_30700) | This study |
| *P. putida* KT2440 | parental | 6 |
| | Δ*gacS* (PP_1650) | This study |
| | Δ*retS* (PP_4824) | This study |
| *P. putida* IsoF | parental | 7 |
| | Δ*gacS* (PisoF_00466) | This study |
| *V. parahaemolyticus* RIMD 2210633 | parental | 8 |
| | Δ*gacA* (VP1945) | This study |
| *E. cloacae* ATCC 13047 | parental | 9 |
| | Δ*tssM* (ECL_RS07530) | 10 |
| *B. thailandensis* E264 | parental | 11 |
| | Δ*tssM-1* (BTH_I2954) | 2 |
| *L. enzymogenes* C3-1 | parental | 12 |
| | Δ*virD4* (GLE_2798) | This study |
| *E. coli* MG1655 | parental | 13 |
| **Plasmids** | | |
| Plasmid | Utility | Source |
| pDMB003_pEXG2_C3_virD4_ko | *L. enzymogenes virD4* deletion allele | This study |
| pDMB005_pEXG2_Pf-5_gacS_ko | *P. protegens gacS* deletion allele | This study |
| pDMB006_pEXG2_KT2440_gacS_ko | *P. putida* KT2440 Δ*gacS* deletion allele | This study |
| pDMB009_pEXG2_Pf-5_retS_ko | *P. protegens* Δ*retS* deletion allele | This study |
| pDMB010_pEXG2_KT2440_retS_ko | *P. putida* KT2440 Δ*retS* deletion allele | This study |

| | | |
|---|---|---|
| pDMB015_pRE112_Vp_gacA_ko | *V. parahaemolyticus* Δ*gacA* deletion allele | This study |
| pDMB038_pEXG2_Pf5_OBC4_ko | *P. protegens* ΔOBC4 deletion allele | This study |
| pDMB041_pEXG2_2-79_gacS_ko | *P. fluorescens* Δ*gacS* deletion allele | This study |
| pDMB043_pEXG2_IsoF_gacS_ko | *P. putida* IsoF Δ*gacS* deletion allele | This study |
| pDMB055_pEXG2_Pf5_phlD_ko | *P. protegens* Δ*phlD* deletion allele | This study |
| pDMB080_pEXG2_Pf5_PARIS_ko | *P. protegens* ΔPARIS deletion allele | This study |
| pDMB082_pEXG2_Pf5_BstA_ko | *P. protegens* Δ*bstA* deletion allele | This study |
| pDMB090_pUC18T-miniTn7T-araC-pBad-PARIS-araE | *P. protegens* PARIS complementation | This study |
| pDMB093_pUC18T-miniTn7T-araC-pBad-aba_bstA-araE | *P. protegens bstA* complementation | This study |
| pEXG2 | Allelic exchange vector | 14 |
| pRK2013 | tri-parental mating helper plasmid | 15 |
| pTNS3 | tri-parental mating helper plasmid | 16 |
| pUC18T-miniTn7T-araC-pBad-PARIS-araE | Arabinose-inducible expression from attTn7 neutral site | 17 |
